# Exogenous application of dsRNA for the control of viruses in cucurbits

**DOI:** 10.1101/2022.03.14.483310

**Authors:** Josemaría Delgado-Martín, Leticia Ruiz, Dirk Janssen, Leonardo Velasco

**Author notes:** **Correspondence:** Leonardo Velasco.

## Abstract

The recurrent emergence of virus diseases in intensive horticultural crops requires alternative control strategies. Topical application of dsRNA molecules homologous to pathogens has been proposed as a tool for plant virus control. These dsRNAs induce the silencing mechanism (RNAi) that degrades homologous dsRNAs. Cucumber green mottle mosaic virus (CGMMV) represents a serious threat to cucurbit crops. Since genetic resistance to the virus is not yet available in commercial varieties, we aimed to control this virus by RNAi. For this purpose, we obtained constructions both for expressing dsRNA in bacteria to treat cucumber plants by topical application and for agroinoculation in experiments done in the growth chamber. Besides, greenhouse tests were performed in spring and in summer when plants were challenged with the virus and differences in several parameters were investigated, including severity of symptoms, dry weight, total height, virus accumulation and virus-derived small interfering RNAs (vsiRNAs). Spraying of plants with dsRNA reduced significatively CGMMV symptoms in the plants in growth chamber tests. Agroinfiltration experiments done under identical conditions were also effective in limiting the progress of CGMMV disease. In the greenhouse assay performed in spring, symptoms were significatively reduced in dsRNA-sprayed plants and the development of the plants improved with respect to non-treated plants. Virus titters and vsiRNAs were clearly reduced in dsRNA-treated plants. The effect of protection of the dsRNA was less evident in the greenhouse assay carried out in summer. Besides, we investigated the mobility of long (ds)RNA derived from spraying or agroinfiltrated dsRNA and found that it could be detected in local, close distal and far distal points from the site of application. VsiRNAs were also detected in local and distal points and the differences in accumulation were compared. In parallel, we investigated the capacity of dsRNAs derived from genes of tomato leaf curl New Delhi virus (ToLCNDV), another economically important virus in cucurbits, to limit the disease in zucchini, both by agroinfiltration or direct spraying, but found no protection effect. In view of the results, topical application of dsRNAs is postulated as a promising strategy for CGMMV control in cucumber.

## 1 Introduction

Food production faces a dual challenge: to increase by 60 % in order to feed the estimated global human population of 10 billion people expected by 2050, while reducing significantly the overall loss of food due to pests and pathogens which now range between 17% and 30% depending on the crop species (FAO, 2019; Savary et al. 2019). Current approaches to the management of these pathogens and diseases are based on the use of chemicals, insecticides and fungicides or on the development of genetic resistances to diseases and/or on the generation of transgenic cultivars. Legislation and consumer demands urge the use of sustainable pest and disease control, and the development of alternatives to improve crop yields as the pressure on global nutrition grows every year (Cagliari et al. 2019). In particular, the increasing number of emerging viral diseases require alternative control strategies (Velasco et al. 2020). As one of such recently developed plant protection tools, the exogenous application of dsRNA to induce RNA interference (RNAi) is one key step-change that can impact the way we will protect crops from pathogen attack in the future (Tenllado et al., 2004; Mitter et al., 2017b; Niehl et al., 2018; Fletcher et al., 2020). RNAi refers to a natural regulatory mechanism in gene expression. This mechanism was discovered in *Caenorhabditis elegans* in 1998 and since then great advances have been made in its potential applications for the control of plant pathogens (Fire et al., 1998; Pugsley et al. 2021). It is mediated mainly by two types of molecules, small-interfering RNAs (siRNAs) and microRNAs (miRNAs) (Sanan-Mishra eta al., 2021). Whereas the former are endogenously derived and involved in the regulation of gene expression, siRNAs can be of exogenous origin from viruses or artificial supply (Matranga and Zamore, 2007), or endogenous derived from transposons (Golden; Gerbase; Sontheimer, 2008). The presence of exogenous dsRNA (double-stranded RNA) generates the activation of DICER and RISC complexes that cleave dsRNA and use the siRNAs as as a template for the degradation of complementary RNAs (Vargason et al. 2003, Pantaleo et al. 2007). This mechanism is known as PTGS (post-transcriptional gene silencing) in plants and is the way plants cope with viral genomes, either by inducing immunity or recovery from infection, depending on whether PTGS occurs before or after virus infection (Pooggin et al, 2003; Vaucheret et al., 2001). The major challenge of RNAi application in plants is to cross the cell wall and reach the cell interior. Several authors have evaluated the effectiveness of topical application of free dsRNA reaching a high effectiveness and a protection time frame of 5-7 days (Tenllado et al. 2003). Others propose the use of nanoparticles (NPs) to improve the delivery conditions of dsRNA to the cellular interior and to release dsRNA in a controlled manner and achieve protection against the virus of 20 days (Mitter et al. 2017a).

Cucumber green mottle mosaic virus (CGMMV) belongs to the genus *Tobamovirus*, family *Virgaviridae* (Adams et al., 2012). It was described as the first-known tobamovirus that infects cucurbits, by Ainsworth in 1935, as a severe threat for cucumber (Ainsworth, 1935). It causes severe symptoms including leaf mottle mosaic, leaf blistering, stunted growth and distortion in fruits and leaves in cucumber but remains symptomless in zucchini (*Cucurbita pepo* L.) (Crespo et al., 2017; Crespo et al., 2019). CGMMV has a 6.4 kb single-stranded, positive-sense RNA genome encapsidated within ∼2,000 molecules of a single species of capsid protein particles, forming rigid rods ca 300 × 18 nm with a helical structure (Hollings et al., 1975). The genomic RNA contains four open reading frames (ORFs) that encode four defined proteins. Two polypeptides are necessary for its replication complex. First, a 129 kDa polypeptide containing methyltransferase and helicase motifs required for RNA replication. Second, a long 186-kDa polypeptide formed by suppression of a UAG termination codon encodes an RNA-dependent RNA polymerase at its carboxyl terminal domain (Crespo et al., 2017). Two additional proteins are translated from subgenomic mRNAs corresponding to ORFs in the 3′ half of the genomic RNA (Dombrovsky et al., 2017). It is seed and mechanically transmitted and is characterized by its stability and ability to persist for long periods without a host. CGMMV is responsible for extensive damages in cucurbit crops. CGMMV is responsible for extensive damages in cucurbit crops, including Spain as one of the leading producers in Europe (Crespo et al. 2017). CGMMV causes in cucumber systemic symptoms such as mottling, mosaic, blistering, leaf stunting and dwarfing (Mandal et al. 2008). No effective genetic resistance to CGMMV is yet available in commercial cucurbits, and the control of the virus consists in limiting the spread in the field thought management practices, such as removing infected material and solarization of the soil, and use of healthy plant material and seeds.

Tomato leaf curl New Delhi virus (ToLCNDV) is a ssDNA virus with a bipartite genome that belongs to genus *Begomovirus*. It has two genome components, named DNA-A and DNA-B (Padidam et al., 1996). DNA-A contains the AV1 (encoding a coat protein) and AV2 genes in the virion sense orientation, and AC1 (coding a virus replication-associated protein), AC2 (encoding a replication enhancer protein), AC3 (coding a transcriptional activator protein) and AC4 in the complementary sense orientation (Padidam et al., 1996). ToLCNDV DNA-B contains the BV1 gene (coding a nuclear shuttle protein) in the virion sense orientation, and the BC1 gene (coding a movement protein) in the complementary sense orientation (Fondong, 2013). The virus is transmitted by the whitefly *Bemisia tabaci* and affects crops from the *Solanaceae* and *Cucurbitaceae* families, causing severe economic loses worldwide. This virus is mainly present in Asia, but recently, a recombinant has spread to the Mediterranean basin, affecting mainly zucchini, cucumber and melon and has tomato as a reservoir host (Janssen et al., 2022; Fortes et al., 2016). In Spain, ToLCNDV infection produces severe symptoms in cucurbits, such as curling and distortion of leaves, green and yellow mosaics and other deformations as well as stunting of the plant. Control of the virus has been achieved by controlling the vector and progress is being made in the search of resistance sources for breeding commercial cultivars (Téllez et al., 2017; Sáez et al., 2016). In this study we evaluated the effect of the exogenous application of homologous dsRNA derived from CGMMV and ToLCNDV in cucumber and zucchini, respectively, to elicits natural RNAi response and eventually resistance to these diseases.

## 2 Materials and Methods

### 2.1. Nucleic acid extraction and the obtention of constructions for the generation of dsRNAs

Total RNA was extracted from young cucumber leaves infected with the Asian isolate of CGMMV (CGSPCu16) detected in Spain (Crespo et al., 2017), using the Spectrum Plant Total RNA Kit (Sigma, Spain) and following the manufacturer’s instructions. Synthesis of cDNA by reverse transcription was performed in 20 μL reactions with 200 ng of total RNA with the High-Capacity cDNA reverse transcription kit (Applied Biosystems, USA). For the subsequent PCRs, primer pairs that included attB adaptors were designed based on the *cp* (coat protein gene) and *mp* (movement protein gene) sequences of CGMMV CGSPCu16 (Supp. Table S1; Delgado-Martin & Velasco, 2021). PCR reactions were carried out with the AmpliTools master mix (Biotools, Spain) using 50 pmol each of forward and reverse primers in the following conditions: an initial denaturing cycle of 2 min at 94 °C, then 40 cycles of 30 s at 95 °C, 30 s at 55 °C and 40 s at 72 °C, and a final extension step of 5 min at 72 °C. The amplicons were cloned into vector pDONR221 using the BP clonase (Thermo Scientific, Spain) and *Escherichia coli* Top10 cells were transformed and selected in LB supplemented with kanamycin. The plasmids obtained, pENT-CP and pENT-MP, were sequenced in order to check the correctness of the insertions. Next, the CP or MP fragments were cloned into vector L4440gtwy that includes attR sites flanked by T7 promoters (G. Caldwell, Addgene plasmid # 11,344) using LR clonase (Thermo Scientific, Spain). The obtained plasmids pL4440-CP and pL4440-MP were used to transform *E. coli* HT115(DE3) cells that were selected in LB supplemented with ampicillin. *E. coli* HT115(DE3) contains in its chromosome the T7 RNA polymerase gene under the *lac* promoter and lacks RNAse III activity. For the constructions derived from ToLCNDV, we used a plant infected with the infective clone of ToLCNDV-ES (Ruiz et al., 2017). Total DNA was extracted from the plant with the GeneJet Plant Genomic DNA purification kit (Thermo Scientific, Spain) and used to produce amplicons from the genes *AV1* and *BC1*, flanked by attB sites (Supp. Table 1). The amplicons were used to obtain plasmids L4440-AV1 and L4440-BC1 following a similar strategy to that described above. These plasmids were used to transform *E. coli* HT115 for the synthesis of the AV1- and BC1-dsRNAs.

### 2.2. *In vivo* production of dsRNAs in *E. coli* HT115

*E*.*coli* HT115(DE3) cells that contained plasmids pL4440-CP, -MP, -AV1 or -BC1 were grown in LB supplemented with ampicillin and IPTG used as inducer of the expression of the T7 RNA polymerase gene. For dsRNA production, 4 mL of the bacteria were grown overnight at 37ºC in LB plus ampicillin. One mL of the culture was used to inoculate 250 mL flasks that contained 100 mL LB, ampicillin (100 μg/mL) and IPTG (1 mM). The cells were grown for about six hours at 37 ºC, when OD determined the steady state of the growth. The dsRNA was isolated from *E. coli* cells using phenolic extractions. Briefly, *E. coli* cells were collected after centrifugation in Falcon tubes at 8,000 x *g*, then resuspended using the vortex in 4 mL of Trisure (Bioline Iberia, Spain) and the lysis was performed after incubation for 5 min at room temperature. Next, 0.2 mL of chloroform was added and mixed up and for phase separation, samples were centrifuged in cold at 12,000 x *g* for 15 min. The upper phase was collected in new tubes and the nucleic acids were precipitated with cold isopropanol. Afterwards, the nucleic acids were resuspended in MilliQ water and examined in 1% agarose gels stained with RedSafe (iNtRON Biotechnology, South-Korea) under UV light.

### 2.3. Constructs for agroinoculation

Plasmid pHellsgate 8 (Helliwell and Waterhouse, 2003) was the backbone vector for cloning the virus segments in direct and reverse directions separated by a hairpin. The plasmid was digested with *Xho*I and *Xba*I (New England Biolabs, UK). Next, amplicons were obtained using the cDNA from CGMMV and ToLCNDV and the primers described in Supp. Table 2. The amplicons, derived from the *cp* and *mp* genes of CGMMV and the *AV1* and *BC1* genes from ToLCNDV were used for Gibson assembly to pHellsgate 8. For each construction, four segments were assembled following manufacturer’s recommendations (New England Biolabs, UK), including the virus gene(s) with direct and reverse positions and the corresponding segments for the *Xba*I-*Xho*I digestions of pHellsgate8 vector that included the hairpin. The resulting plasmids, pGHE-CP, pGHE-MP, pGHE-AV1 and pGHE-BC1 were used to transform Top10 *E. coli* cells that were grown in LB plates supplemented with streptomycin (100 μg/ml). Plasmids were extracted from the cells and checked by restriction fragment analysis and Sanger sequencing. The plasmids were then transferred to *Agrobacterium tumefaciens* LBA4404 by electroporation and selected with the same antibiotic.

### 2.4. Agroinoculation procedures

For the agroinoculation with the *A. tumefaciens* LBA4404 strain carrying either pGHE-CP/pGHE-MP in cucumber or pGHE-AV1/pGHE-BC1 in zucchini, one colony of each clone was collected in 5 mL of liquid LB and streptomycin and grown overnight at 30°C with constant shaking. Next, the cells were harvested by centrifugation at 3,000 x *g* for 10 min at 4°C and resuspend in infiltration medium (10 mM MES pH 5.8, 200 μM acetosyringone) adjusting the OD_600_ to 0.5. The agroinoculation in the abaxial side of fully expanded leaves and/or cotyledons of the plants was performed using a needleless 2 mL syringe. Virus infection and symptom inspection was performed as described above.

### 2.5. Plant material, inoculum source and data analyses

*Cucumis sativus* cv ‘Bellpuig’ (Fitó, Spain) was used in all CGMMV experiments. In growth chamber experiments (phytotron), the plants were treated with the CP/MP-dsRNAs using several different approaches at two true fully expanded leaves. In two separate greenhouse experiments, one performed in the month of April, and a second in July the of 2021, plants were treated with the CP-dsRNA at three fully expanded leaves. The dsRNA was resuspended in water and applied either by rubbing on two contiguous leaves using of 60 μg of total RNA/leaf or alternatively by spraying the same amount of total RNA using an artist airbrush at 2.5 bar pressure. After the dsRNA treatments, the virus was mechanically inoculated in the cotyledons as described elsewhere (Crespo et al., 2018). For the virus challenge, 1 g of leaf material from a *C. sativus* plant infected with the Asian isolate of CGMMV (CGSPCu16) was used as inoculum source (Crespo et al., 2017). The plant tissue was crushed in cold with a pestle in the mortar in phosphate buffer (5 mM, pH 7) and activated charcoal. Prior to inoculation, carborundum powder was applied on the cotyledons. At 18-25 days post inoculation (dpi), symptoms were recorded and rated (Supp. Fig. 1), as well as the total length of the plants. Dry weight of the complete plants was registered for comparisons. Samples were taken for RNA extraction and virus and virus-derived small interfering RNAs (vsiRNAs) quantitation. In the greenhouse experiments, three groups of fifteen plants were used in each experimental condition: mock-treated plants, non-inoculated plants and dsRNA-treated plants. The effect of virus-species-specific dsRNAs on plants infected with ToLCNDV was studied in groups of ten zucchini (*Cucurbita pepo* cv. ‘Victoria’, Clause, Spain) plants. Treatments consisted of rubbing with 120 μg each of the AV1- and BC1-dsRNA. Infection was done using viruliferous whiteflies reared on plants previously infected with ToLCNDV-ES. Another group of ten plants was inoculated but not treated and used as a control. Plants were kept in the growth chamber and symptoms were inspected at 18 dpi. The statistical analyses of the data were conducted by ANOVA followed by mean separation using the post hoc Tukey (equal variances) test as available in JAMOVI v.2.2.3 statistical suite (Jamovi Project, 2020: https://www.jamovi.org).

### 2.6. Small RNA sequencing and analysis

RNA extracts of CP-dsRNA-treated and mock (non-treated) plants from the greenhouse assay performed in spring, were pooled in two independent samples (DS and MO, respectively). The small RNA fractions were excised from gels and the cDNA libraries were prepared for High-throughput sequencing (HTS) as single reads with the Illumina platform using the services provided by Sistemas Genómicos (Valencia, Spain). The Illumina sequencing adapter was trimmed off from the raw sequences and reads between 18 and 24 nt length were used in the subsequent analyses. The small RNA populations were aligned to the indexed CGMMV CGSPCu16 genomic sequence using the Bowtie2 module (Langmead & Salzberg, 2012) as available in Geneious (Biomatters). The BAM alignment file produced was processed using MISIS-2 (Seguin et al., 2014) and the tables generated were graphically represented in Veusz (Sanders, 2021: https://veusz.github.io/).

### 2.7. CGMMV quantitation by RT-qPCR

A hole punch was used to obtain approximately 100 mg of leaf tissue from each sample (15 biological replicates per condition) and the total RNAs were extracted using the Spectrum Plant RNA kit (Merck, Spain). RNAs were quantified using the NanoDrop ND-1000 (Thermo Fisher Scientific, Waltham, MA, USA). Subsequently, cDNA was produced using 2 μg of the total RNA extract from each plant and the High-Capacity cDNA Reverse Transcription in 20 μl reaction volume and following the manufacturers’ instructions. Each qPCR reaction (20 μl final volume), in triplicate, contained 1 μl of the cDNA, 10 μl of KAPA SYBR Green qPCR mix (KAPA Biosystems, MA, USA), and 500 nM of each CGMMV *cp* or *mp* primers (Supp. Table 2). In separate reactions, we included the primers for the *C. sativus* 18S rRNA gene as a reference. Specificity of the amplicons obtained was checked with the Bio-Rad Optical System Software v.2.1 by means of melting-curve analyses (60 s at 95°C and 60 s at 55°C), followed by fluorescence measurements (from 55–95°C, with increments by 0.5°C). The geometric mean of their expression ratios was used as the normalization factor in all samples for measuring the quantification cycle (Cq). The relative expressions of the CGMMV amounts were calculated based on the comparative Cq (2^−ΔΔCq^) method as described by Livak and Schmittgen (2001).

### 2.8. Quantitation of vsiRNAs

CGMMV vsiRNAs were quantified by RT-qPCR according to Shi and Chiang (2005) with some modifications: 2 μg of RNA extracts from the cucumber leaves were treated with DNaseI (Merck, Spain) and polyadenylated using the Poly(A) polymerase (New England Biolabs, UK)) and ATP, following the manufacturer’s recommendations. Next, the polyadenylated RNA was precipitated with ethanol, NaOAc 3M, pH 5.2 and glycogen (Merck, Spain) in cold, resuspended and reverse transcribed with the High-Capacity cDNA Reverse Transcription Kit and 0.25 μM of the poly(T) adapter (Supp. Table 1). The amplifications were carried out with specific primers, designed upon the vsiRNA hotspots for the CGMMV *RdRp, mp* and *cp* genes and the universal 3′-adapter reverse primer (URP). A primer based on the 5.8S rRNA was designed as a reference in the amplifications. Each reaction (20 μL final volume) contained 1 μL of the diluted cDNA, 10 μL of KAPA SYBR Green qPCR mix (KAPA Biosystems, USA) and 500 nM each of the specific and the URP primers. The cycling conditions consisted of an initial denaturation at 95 °C for 10 min, followed by 45 cycles at 95 °C for 15 s, 58 °C for 20 s and 60 °C for 40 s. Five biological repetitions were included in each case. Each qPCR (biological sample), including those for the 5.8S as internal control, was repeated three times. The specificity of the amplicons obtained was checked as above. The relative expressions of the vsiRNAs were calculated based on the comparative Cq method as above.

## 3 Results

### 3.1. Generation of CGMMV and ToLCNDV-derived dsRNA

By Gateway cloning of the viral amplicons we obtained L4440-derived plasmids carrying partial segments of the *AV1* and *BC1* of ToLCNDV and the *cp* or *mp* gene of CGMMV flanked by two IPTG-inducible T7 promoters. These plasmids were subsequently introduced by transformation into the RNAse III-deficient strain *Escherichia coli* HT115(DE3) for dsRNA expression. To analyze dsRNA synthesis from HT115 cells we extracted total RNA by the Trisure method after 7 h of incubation with IPTG as inducer. DsRNA of the expected sizes, 720, 700, 590, and 650 bp, were obtained in HT115 cells harboring plasmids L4440-AV1, -BC1, -CP or -MP, respectively, when induced with IPTG (Supp. Fig. 2). Yields were ∼120 μg of total RNA extract per 5 ml of bacterial culture, which will allow us to treat a plant with 60 μg per leaf, the standard amount used in the assays described here.

### 3.2. Phenotypic effects of dsRNA applications on CGMMV-inoculated cucumber plants

Preliminary experiments were performed in the phytotron in order to test the effect of CGMMV-derived dsRNAs, either in the form of rubbing, spraying or agroinoculation. In the initial experiment, five cucumber plants at cotyledon stage were used in each of the following treatments mock and CP/MP-dsRNA treatment by rubbing. The same day, the plants were inoculated (challenge) with CGSPCu16. At 15 dpi, 2 mock inoculated plants initially showed virus symptoms that finally appeared in all the mock-inoculated plants at 18 dpi. In contrast, the dsRNA-treated plants remained symptomless in this period (Fig. 1). In a second experiment, performed in the phytotron, 10 plants in each condition were used. In this case, the virus challenge was done three days after the dsRNA application. At 25 dpi, symptoms in two dsRNA-treated plant rated severe (2), while seven remained asymptomatic and one resulted dead. Among the mock-inoculated plants, four resulted dead and three showed very severe symptoms (rate = 3). However, three mock-treated CGMMV-inoculated plants remained asymptomatic in this period (Table 1). Other preliminary experiments showed that spraying with dsRNA was as effective as rubbing in limiting CGMMV disease severity (not shown). An additional experiment was performed in the phytotron to trigger dsRNA synthesis in the plant by agroinoculation with the pGHE-CP and pGHE-MP constructions mediated by *A. tumefaciens*. A group of 10 plants was agroinfiltrated at 2-3 true leaves stage, while a second group of 10 plants was mock-treated. Both groups were challenged subsequently with the virus on the same day. The results showed that the agroinoculation limited the disease progress and protected the plants against CGMMV (Table 2). At 13 dpi all the mock treated plants displayed symptoms. Al 15 days post treatment (dpt)/dpi, four of the agroinfiltrated plants showed symptoms while the rest remained asymptomatic. Finally, at 30 dpt/dpi, eight agroinfiltrated plants showed severe symptoms, one remained asymptomatic and another one showed mild symptoms.

**Fig. 1.**
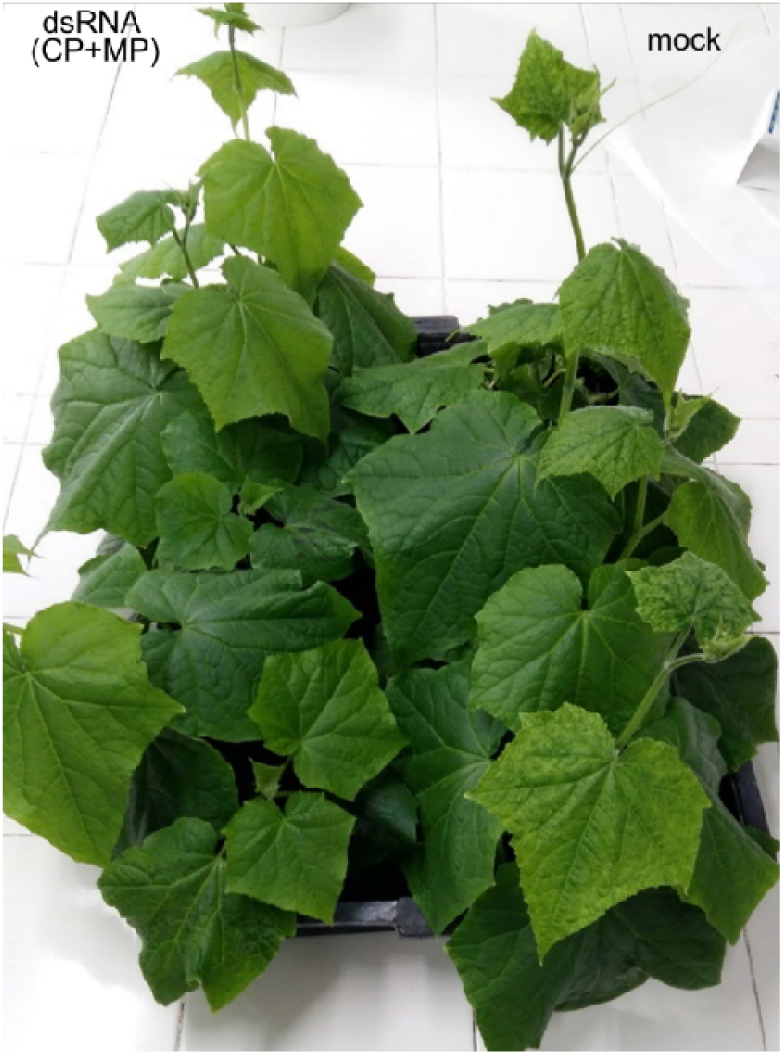
Aspect of the plants inoculated with CGMMV at 25 dpi. Plants were treated prior the inoculations either with mock or with a mix of CP/MP-dsRNAs by rubbing.

**Table. 1.** Effects of CP/MP dsRNA treatments in plants inoculated with CGMMV Sp16 at 25 dpi when grown in the phytotron (second assay). Plants were challenged with the virus three days after the dsRNA application. Parenthesis means dead plants.

| Condition | No symp | Mild symp. | Severe symptoms |
| --- | --- | --- | --- |
| mock | 0/10 | 3/10 | 7(4)/10 |
| CP/MP-dsRNA | 7/10 | 2/10 | 1(1)/10 |

**Table 2.** Occurrence of symptoms with time (days post inoculation, dpi) in plants agroinoculated with pGHE-CP and pGHE-MP mediated by *A. tumefaciens* LBA4404 compared with the mock control plants after challenging with CGMMV Sp16. *One plant remained asymptomatic at 30 dpi and another one showed only mild symptoms. The rest of the plants developed severe of very severe symptoms.

| Condition | Symptoms at: |  |  |  |
| --- | --- | --- | --- | --- |
|  | 15 dpi | 18 dpi | 21 dpi | 30 dpi |
| mock | 10/10 | 10/10 | 10/10 | 10/10 |
| pGHE-CP/pGHE-MP | 4/10 | 5/10 | 7/10 | 9*/10 |

Once the experiments in the phytotron showed an effect of the dsRNAs on limiting CGMMV disease progress, we considered experimental assays under greenhouse conditions (Fig. 2). Two separate experiments were performed. A first experiment was started in mid-April 2021 (23.5 ºC average temperature in the greenhouse), when 15 plants were mock-treated and 15 plants of another group were sprayed with CP-dsRNA at 2-3 leaves stage. Subsequently, all the plants were inoculated with CGSPCu16. A third group of 15 plants was not inoculated as a reference. The treatment was performed by spraying with 60 μg of bacterial dsRNA extract in each of two true leaves per plant (120 μg/plant). At 3 dpt, the plants were challenged with the virus. At 18 dpi (days post inoculation) the cucumber plants were harvested because almost all the mock-treated CGMMV inoculated plants showed severe or very severe symptoms. Then, the dry weight, total height and level of symptom expression were determined (Fig. 3). At this stage samples were taken for RNA extraction and virus quantitation by RT-qPCR. Although we found only small differences between dry weight values, the control non-inoculated plants resulted significantly more robust than those inoculated with CGSPCu16 (*P* = 0.001) but not significatively different with respect to the dsRNA-treated plants (*P* = 0.378). DsRNA treated and mock treated plants slightly, but significantly in dry weight (*P* = 0.036). As for the height, the control and the dsRNA-treated inoculated plants showed no significant differences (*P* = 0.954), in contrast with the dsRNA-untreated inoculated plants, that showed a significantly lower height with respect the control (*P* = 0.005) and the dsRNA treated plants (*P* = 0.005) (Fig. 3). The greatest differences were observed in the expression of disease symptoms. All the mock-treated virus-inoculated plants showed severe symptoms (average rating = 2.13) at 18 dpi. In contrast, four of dsRNA treated plants showed mild symptoms (rating = 1) and the rest remained asymptomatic. No symptoms appeared in the control plants.

**Fig. 2.**
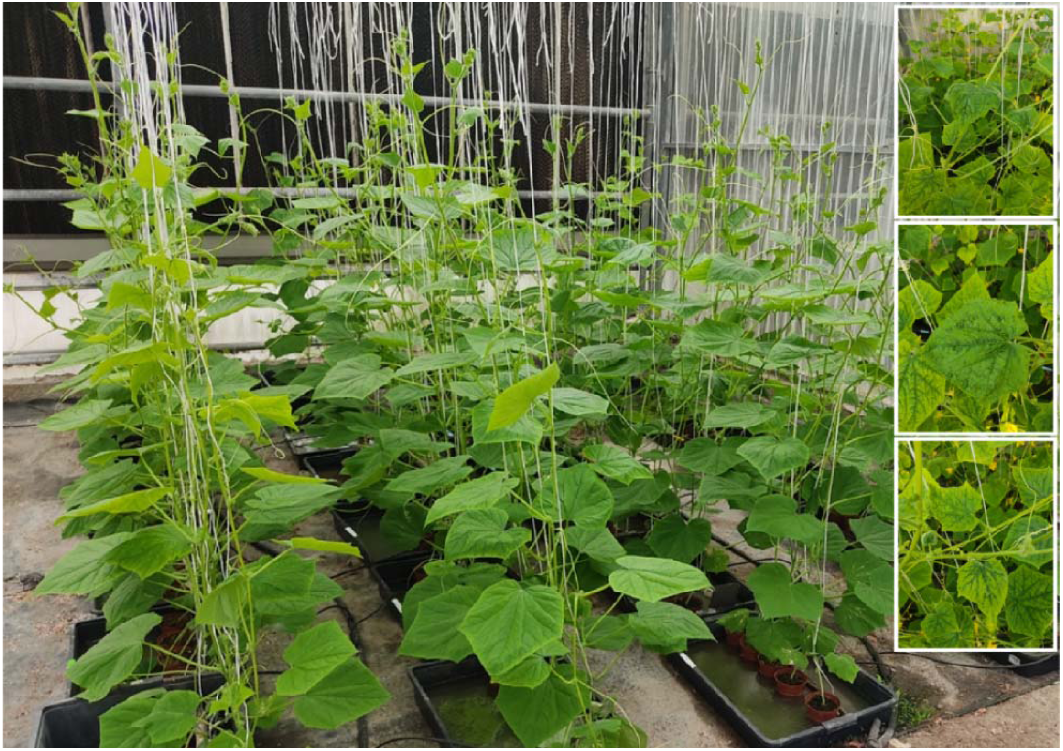
Cucumber plants in the greenhouse assays for the CP-dsRNA treatments and CGMMV inoculation assays. Details of severe virus symptoms are shown in the picture boxes at the right.

**Fig. 3.**
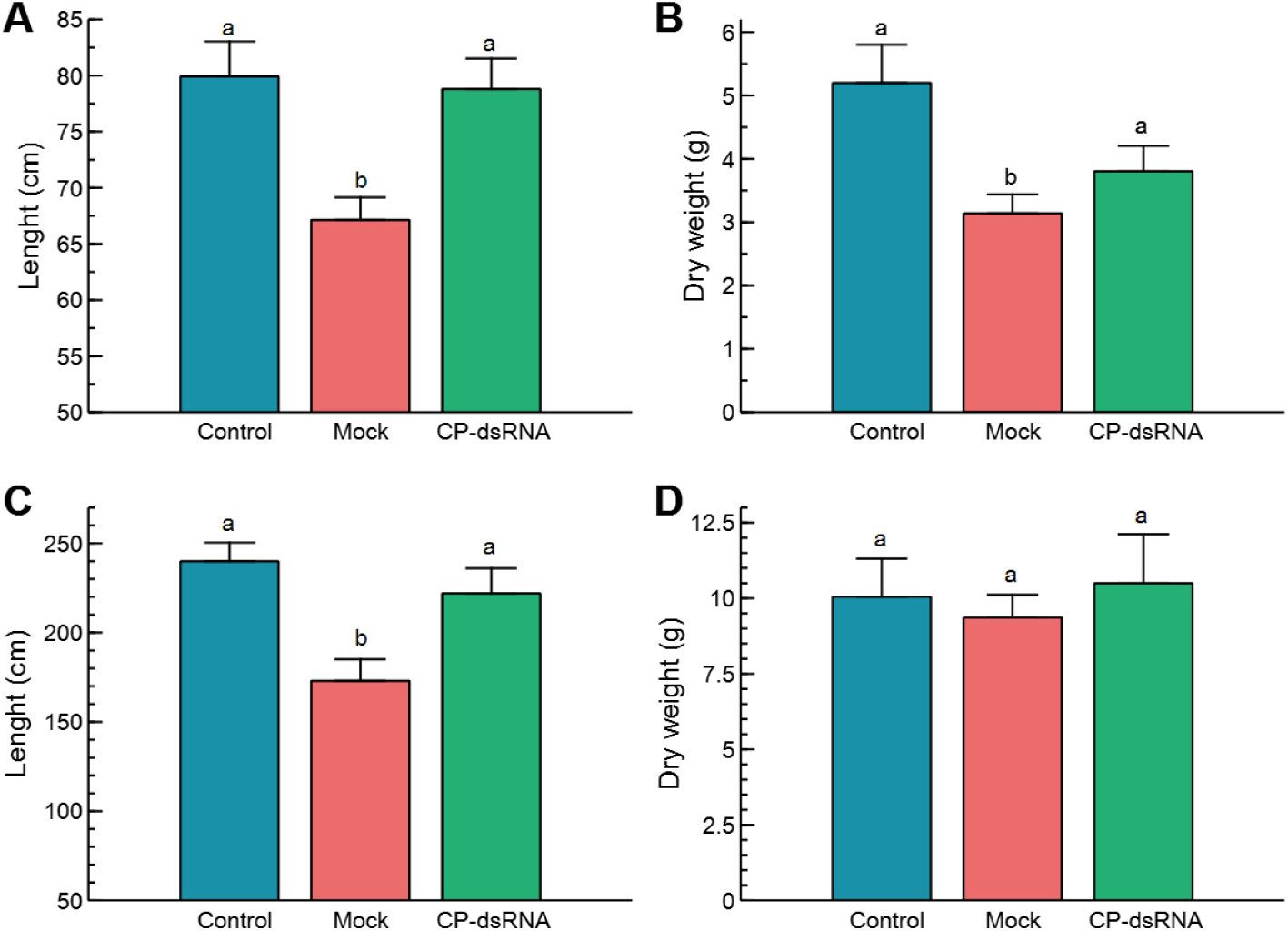
Phenotypical parameters in the plants following CGMMV inoculation and CP-dsRNA treatment. Plants were evaluated at 18 dpi, when the symptoms in all the mock-treated plants rated severe (= 2) or very severe (= 3). Control refers to non-inoculated plants. Bars correspond to the standard error of the means. Different letters on the columns refer to significant differences.

A similar greenhouse experimental approach was performed in summer, at higher temperatures (28 ºC average), starting in the beginning of July 2021. In this case, there were non-significant differences in dry weight among the treatments (*P* = 0.495), but significant differences appeared in length between the dsRNA treated and the non-treated plants (*P* = 0.018). With respect to the control non-inoculated plants, there were no differences with respect to the dsRNA-treated plants (*P* = 0.560). Regarding the differences in length between control non-inoculated and mock-treated plants, the result was significantly different (*P* = 0.001) (Fig. 3). With respect to symptom expression, all the mock-treated plants showed symptoms rating 3 (very severe) at 18 dpi. The dsRNA-treated plants showed an average symptom rate of 1.6, with two plants showing very severe symptoms, one that remained asymptomatic, seven with mild symptoms and other five that showed severe symptoms. Plants in the experiment carried out in July were 70% higher than those grown in April and (dry) weighted 62% more. There is significant correlation (*R*^*2*^) between length and weight, 0.81 and 0.72 for the spring and summer experiments, respectively (Fig. 4). Linear correlation analysis showed differences in the behavior of the treatments between both seasons that showed a greater effect of the dsRNA treatments in limiting disease effects during April (Supp. Table 3).

**Fig. 4.**
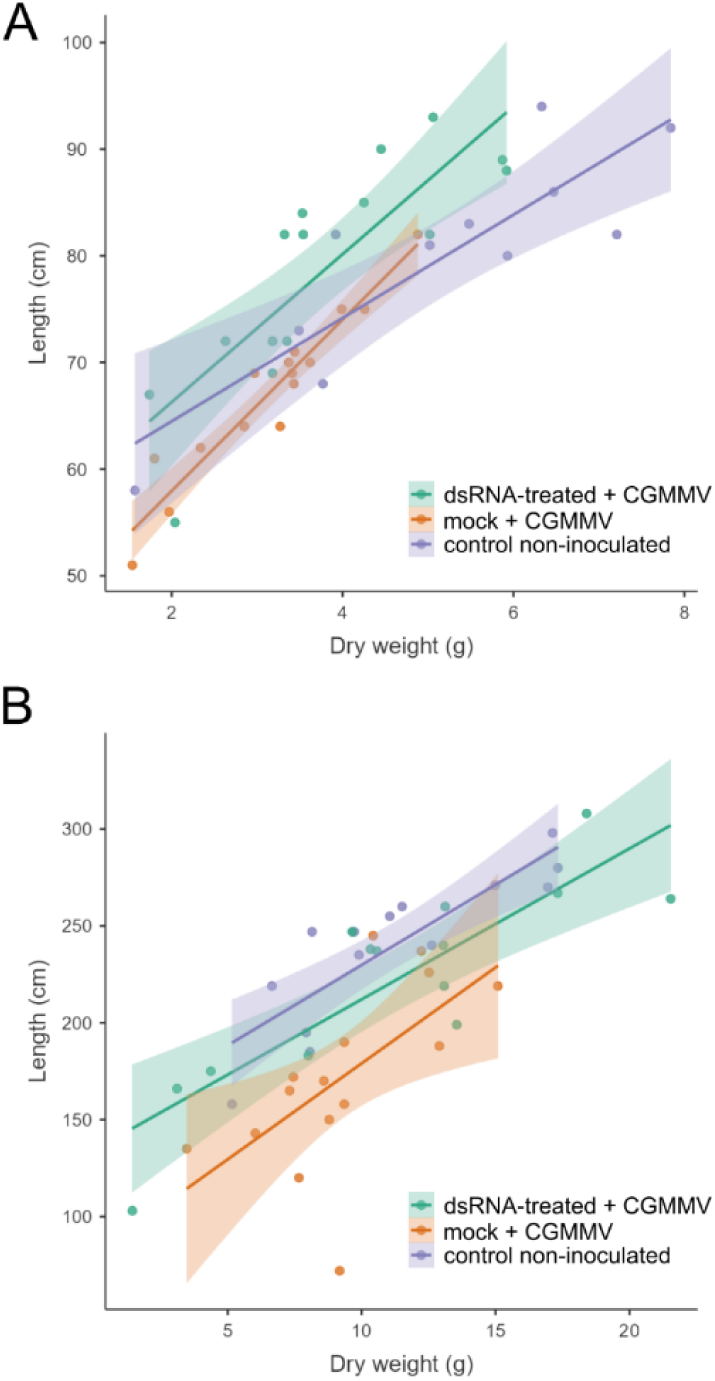
Scatter plot between length and dry weight of the plants of the experiments carried out in spring and summer. For the linear regression analysis see Supp. Table 3.

### 3.3. Quantitation of CGMMV viral RNA in dsRNA-treated and untreated plants

The determinations of relative viral accumulation were performed by comparing the differences in the expression of the CGMMV *cp* gene and the expression of the 18S rRNA gene of *C. sativus* in dsRNA-treated and untreated plants. In the spring assay, at 18 dpi, the average Cq for the *cp* gene in the untreated plants was 15.0 ± 1.8 and for the 18S rRNA the resulting average Cq was 14.3 ± 1.9. In the dsRNA-treated plants the average Cq was 18.7 ± 3.4 for the *cp* and 15.2 ± 2.6 for the 18S Rrna (Fig. 5). Calculation of the ΔΔC_CP-18S_ values resulted in a relative increase of 47.7-fold of virus accumulation in the untreated vs. the dsRNA-treated plants. Expression of the CGMMV *cp* gene was observed in all the inoculated plants, both in the untreated and in the dsRNA treated ones, so that there were no escapes in virus inoculation. In the assay carried out in summer, the difference in ΔCq_CP-18S_ of the untreated compared with the treated plants was less evident than in the spring assay. In this case, the reduction in CGMMV *cp* expression of the dsRNA-treated plants vs. the untreated plants was only 5.8-fold (Fig. 5). All non-inoculated plants tested negative to the *cp* amplification in the RT-qPCRs. When the viral accumulation was compared between the seasons, we found for the untreated plants in summer a 30-fold higher amount with respect to spring. However, when we compared amount of virus in the treated plants, the difference between summer and spring was 243-fold. Thus, the effect of the dsRNA in reducing virus accumulation was significantly higher in spring.

**Fig. 5.**
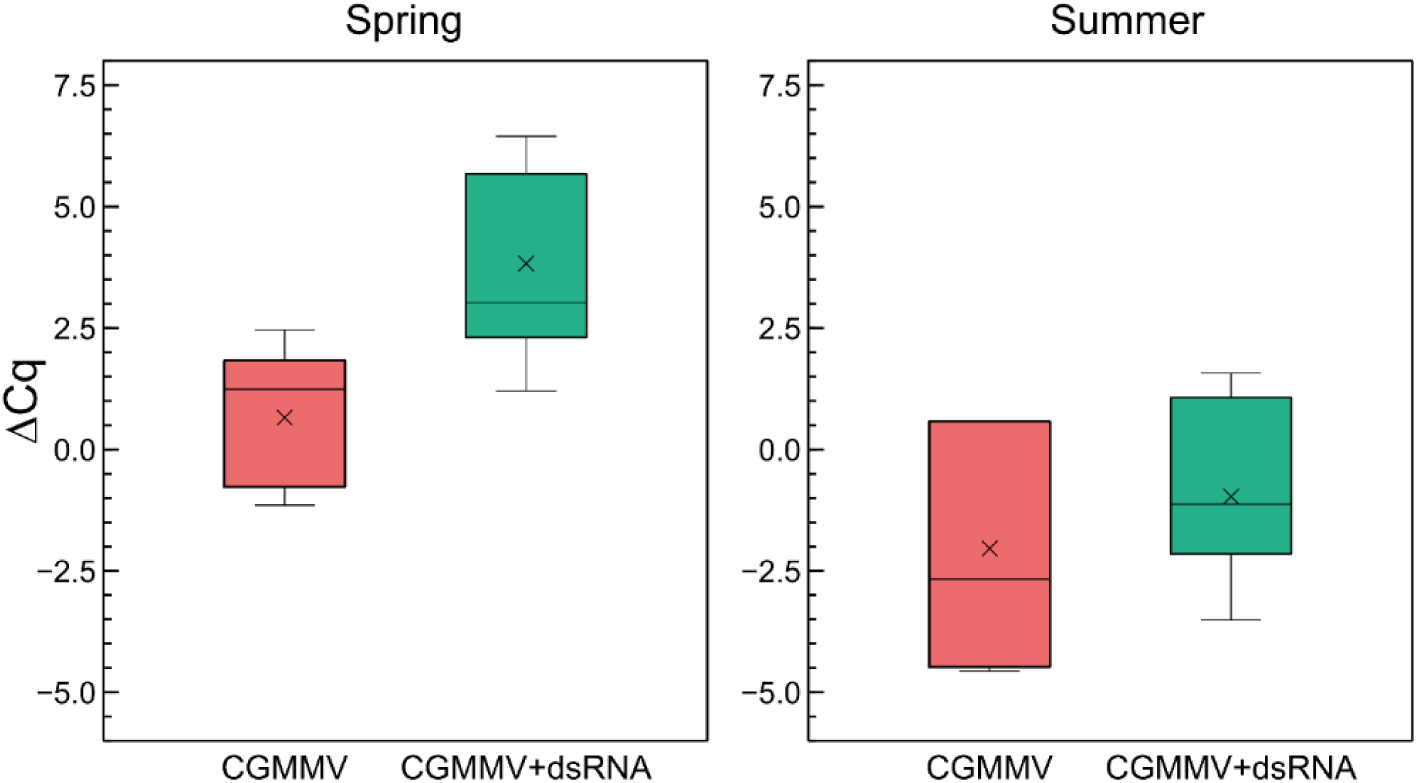
Quantitative differences in CGMMV accumulation between dsRNA-treated and untreated infected plants in two greenhouse assays carried out in spring and summer. The ΔCq refers to the differences between the averages of the Cq of the virus *cp* gene and average Cq of the cucumber internal rRNA 18S control. Bars correspond to the standard deviations.

### 3.4. HTS of vsiRNAs in CGMMV infected plants

From the HTS sequencing of the small RNAs, a total of 12,967,977 raw reads were obtained from the inoculated and dsRNA-treated pool of samples (DS) and 12,819,530 reads from the non-treated inoculated plants (MO). After trimming the adapter and the low-quality reads, 3,162,386 and 6,355,112 reads between 18-24 nt were obtained from each sample, respectively. A profiling of the small RNA reads in function of length resulted when using the *C. sativus* reference in the sRNAtoolbox (Supp. Fig. 3) (Rueda et al., 2019). Read length distributions resulted classified from the analyses according to origin (Supp. Fig. 4). Alignment of the vsiRNAs to the CGMMV genome showed differences between the DS and MO samples. The 21-24 nt reads were aligned to the CGMMV genome (Genbank Acc. No. MH271441), resulting that 199,534 vsiRNAs matched to CGMMV in sample DS and 692,505 vsiRNAs in MO. CGMMV vsiRNAs represented the 12.7 % of total 18-24 nt siRNAs in the MO sample and only the 7.6% in the DS sample (dsRNA-treated plants). Although in both samples, similar hotspots were observed in the *mp* and *cp* genes, a higher prevalence of vsiRNAs matching to the replicase region (*RdRp* gene) was observed in the MO sample (Fig. 6). From the 21-24 nt vsiRNAs aligning to CGMMV, 95% belong to the 21-22 nt class (Supp. Fig. 5). CGMMV vsiRNAs of 21 nt class clearly prevailed (55.06%), followed by the 22-nt class (41.09%) in sample DS. Similarly, the vsiRNAs 21 nt (54.41%) prevailed over the 22-nt class (41.54%) in MO. About 55% of the 21 to 22-nt vsiRNAs aligning to the CGMMV genome were of negative polarity (69.7% in DS; 72.6 in MO), showing a bias towards antisense vsiRNAs. Adenine was the prevailing base at the 5’ end of the vsiRNAs (41.5% and 41.1% for DS and MO, respectively) and uracil at the 3’ end (45.6% and 43.2% for DS and MO, respectively) (Supp. Fig. 6).

**Fig. 6.**
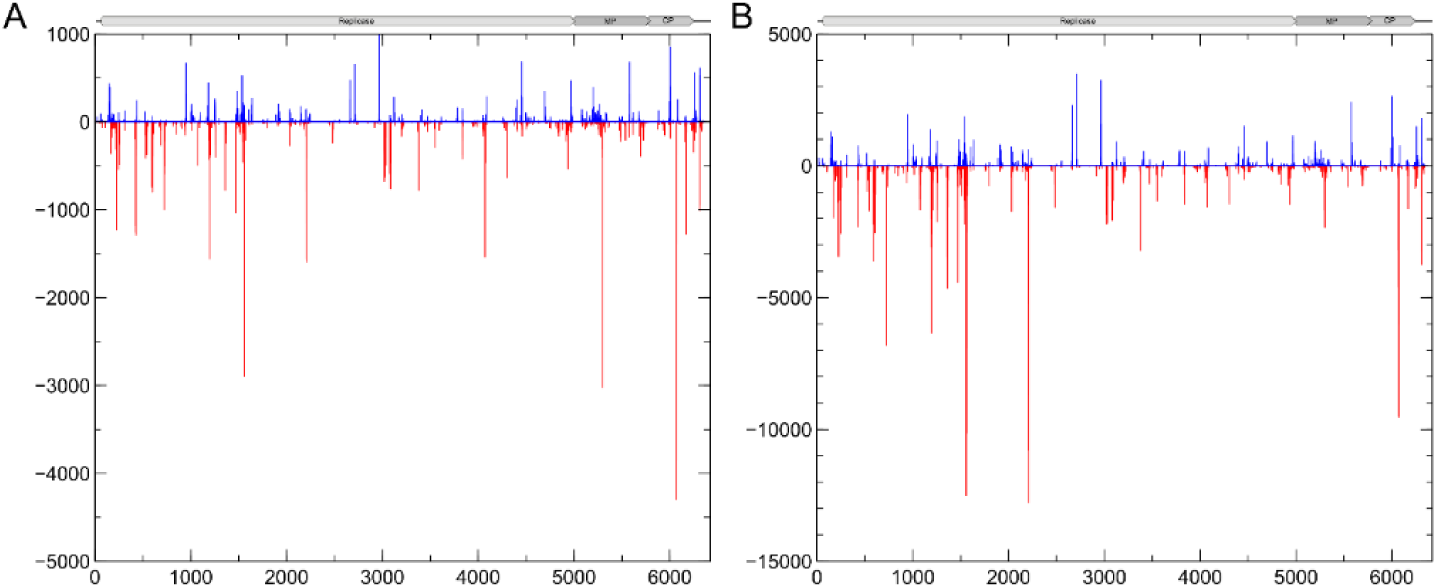
Profile distribution of the small RNAs of the 21-24 nt classes aligning to the genome of CGMMV: (A) Sample DS; (B) sample MO. Peaks above and below the X axis stand for sense and antisense vsiRNA orientations, respectively. A schematic view of the CGMMV genome is represented for reference. Separation between marks in the X axis represents 1 Kb.

### 3.5. RT-qPCR of vsiRNAs in dsRNA-treated and untreated CGMMV infected plants

As the HTS allowed the identification of several hotspots of vsiRNAs aligning to the CGMMV genome (Fig. 6), we could design specific primers for vsiRNA detection and quantitation among plants and conditions (Supp. Table 1). In the assay carried out in spring, CGMMV vsiRNAs were detected in dsRNA treated and non-treated plants at 18 dpi (Figure 7). Non-inoculated control plants showed no specific amplification of the vsiRNAs as shown by the analysis of the melting curves (Supp. Fig. 7). DsRNA-treated plants showed a significant lower amount of specific vsiRNAs than the non-treated plants (Figure 7), being 7.4-fold lower for the 1193-vsiRNA, located in the *RdRp* gene, 14-fold lesser for the 5234-vsiRNA, matching the *mp* gene and 15.8-fold lower for the 6125-vsiRNA, that corresponded to the *cp* gene of the virus. The 6125-vsiRNAs resulted the most abundant in both the dsRNA treated and untreated plants.

**Fig. 7.**
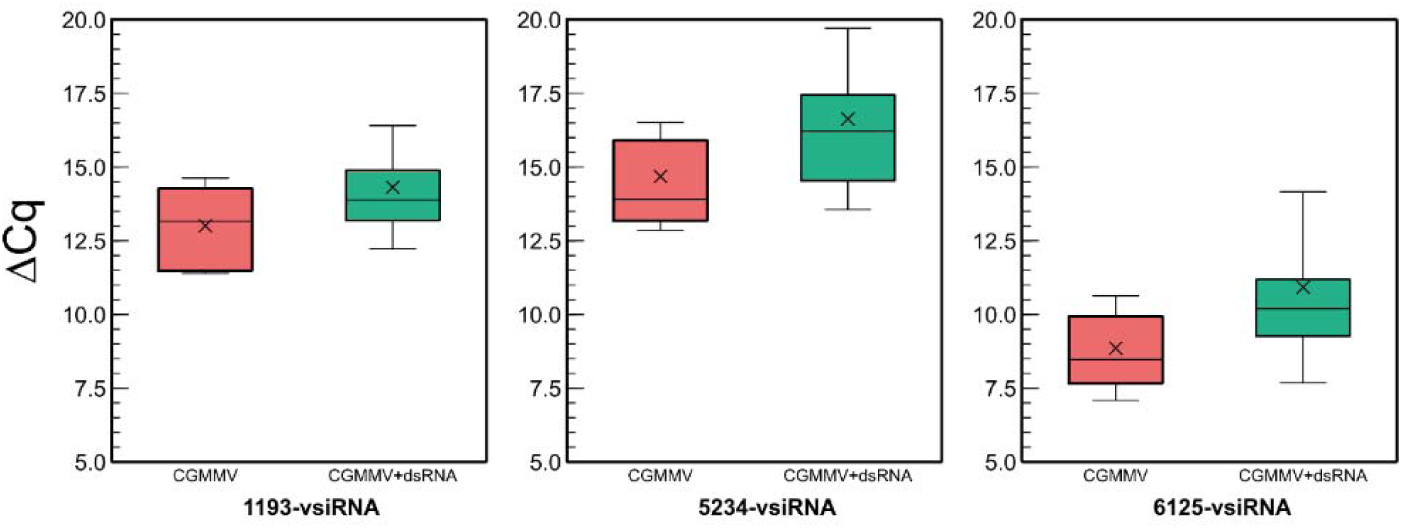
Quantitative differences in vsiRNA titters between untreated (red) and dsRNA-treated (green) CGMMV-infected plants in the experiment carried out in spring. The ΔCq refers to the differences between the Cq of the different vsiRNAs and the cucumber rRNA control (5.8-rRNA). Bars correspond to the standard deviations.

### 3.6. Systemic movements of exogenous CGMMV (ds)RNAs and derived vsiRNAs to distal and proximal parts of the plant

We investigated the systemic movements of long (ds)RNAs and derived siRNAs after the application of dsRNA. Here we define (ds)RNAs as long RNA molecules that move in a cell autonomous (apoplast, phloem) or non-cell autonomous way (symplast), in the form of single or double stranded. In a set of independent experiments performed in the growth chamber, we applied CP-dsRNA by spraying on plant leaves of five plantlets (point 1 in Fig. 8). The leaves were washed thoroughly and we analyzed the presence of (ds)RNAs and vsiRNAs at 3 dpt in distal leaves (point 3), that were foil-covered prior spraying. In a parallel experiment, other five plants per condition were agroinfiltrated with pGHE-CP or pGHE-MP in point 1 (Supp. Fig. 8). Total RNA was then extracted from the sampling point 3 and used for the quantitation of CP- or MP-(ds)RNA and the 6126-vsiRNA or 5324-vsiRNA. Analysis of the results showed that the (ds)RNAs and the vsiRNAs were detected in distal leaves (Fig. 9), both in agroinoculated and in sprayed plants. Comparison of the ΔCq for agroinoculation of MP-dsRNA and CP-dsRNA of the long (ds)RNAs detected in distal leaves showed similar values, but lower to the ones obtained with sprayed dsRNA (Fig. 9A). The calculated ΔΔCq values of leaves that were dsRNA-sprayed and agroinfiltrated differed 9-fold. On the other hand, comparison of ΔΔCq values showed that the amounts of vsiRNAs derived from the CP- and MP-dsRNAs in agroinfiltrated plants were in the same range, while the 6125-vsiRNAs derived from the CP-dsRNA in sprayed plants was 14-fold more abundant (Fig. 9A). With respect to long RNAs, they were slightly more concentrated in distal leaves of plants that were sprayed when compared with those from plants that had been agroinoculated (Fig. 9B).

**Fig. 8.**
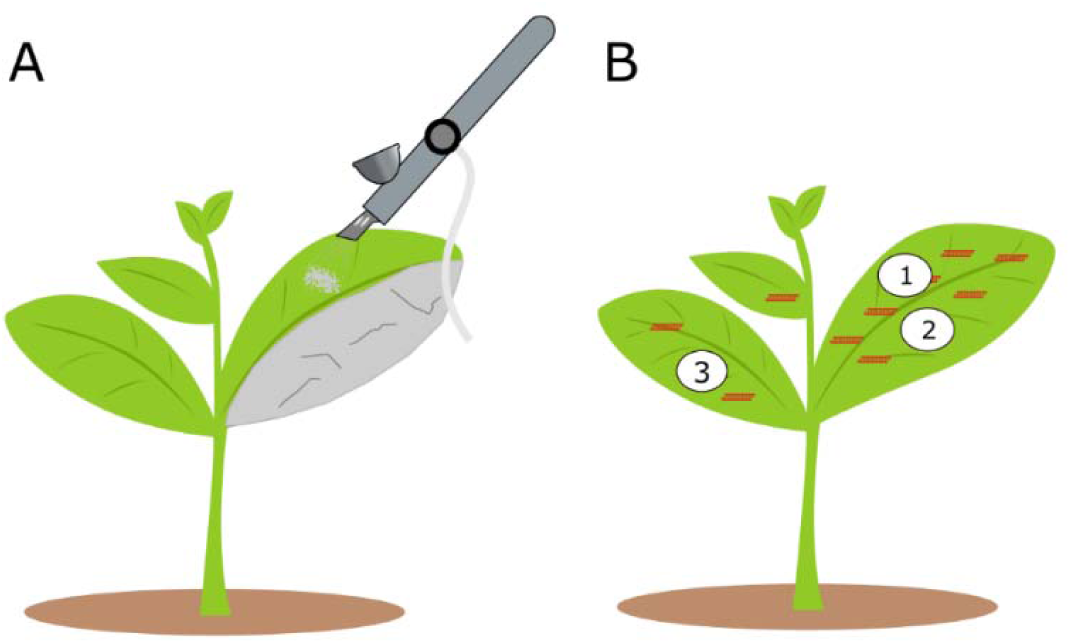
The CP-dsRNA was applied by spraying with the airbrush onto a leaf, one half of which and the rest of the plant, were covered with aluminum foil to prevent direct entry of the dsRNA (A). Points of sampling at three days post application for detecting *cp*-derived RNAs and specific vsiRNAs (B).

**Fig. 9.**
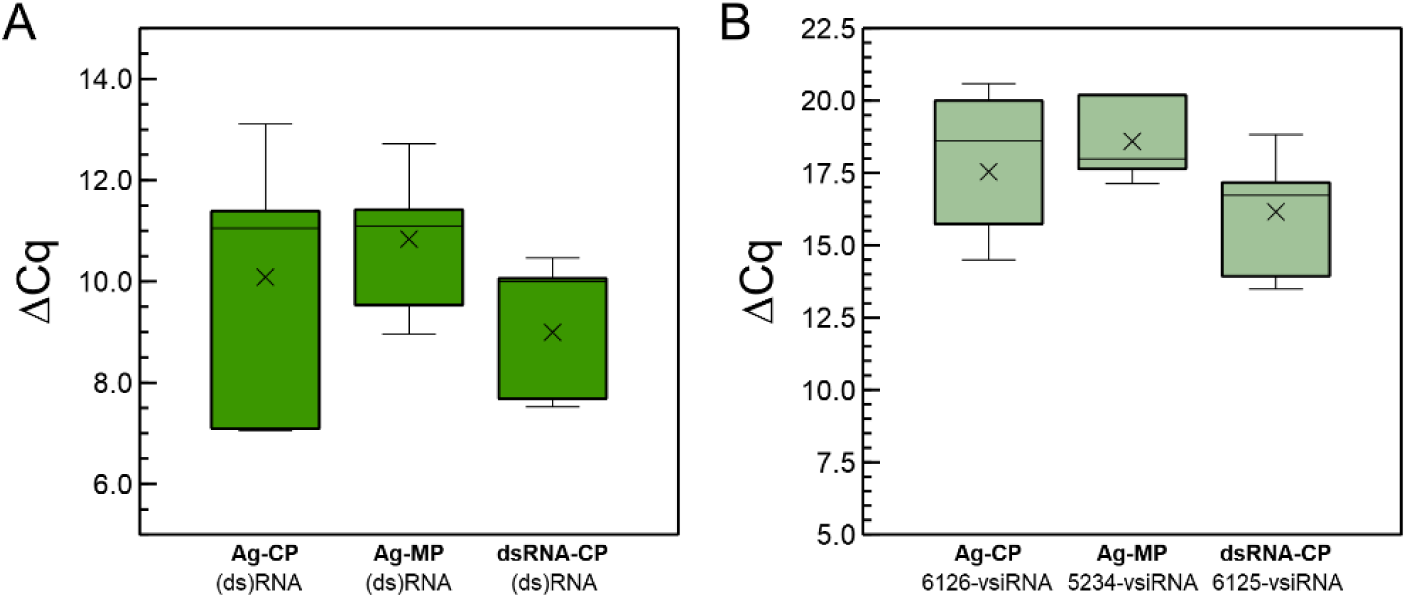
Quantitative differences at 3 dpt in (A) CP-ds(RNA) and (B) 5234- or 6125-vsiRNA amounts in distal (point 2) non-treated leaves among plants either directly sprayed with CP-dsRNA (dsRNA-CP) or agroinfiltrated with the CP-dsRNA (Ag-CP) or the MP-dsRNA (Ag-MP) expressing constructions pGHE-CP and pGHE-MP mediated by *A. tumefaciens* LBA4404.

In another experiment, we included six plants per condition and analyzed two additional points of sampling, including the site of application, to investigate the movement of the long RNAs and derived siRNAs that is close distal to the site of application of the dsRNA (point 2 in Fig. 9). Plants were prepared so that the dsRNAs were applied only on half of the leaves, keeping the other halves and the rest of the plant foil-covered to prevent them to be reached by the sprayed CP-dsRNA (Supp. Fig. 9). Three points were thus evaluated by RT-qPCR (Fig. 8B). Values of ΔCq_CP-18S_ for the long (ds)RNAs are displayed in Fig. 10A. Calculation of the ΔΔCq_CP-18S_ enabled us to establish that that at point 2 and 3, there was 3.8 × 10^4^-fold and 1.55 × 10^5^-fold less CP-(ds)RNA, respectively, with respect to the point of spraying the CP-dsRNA (point 1). Thus, about four orders of magnitude less (ds)RNA moved from the point of application to the close distal part of the leaf, and from there, even a five order of magnitude less long RNA molecules moved to another leaf (far distal). When we compared the differences in CP-(ds)RNA amounts between the close distal (point 2) and the far distal point 3, the difference was only 50-fold of the former with respect to the latter.

**Fig. 10.**
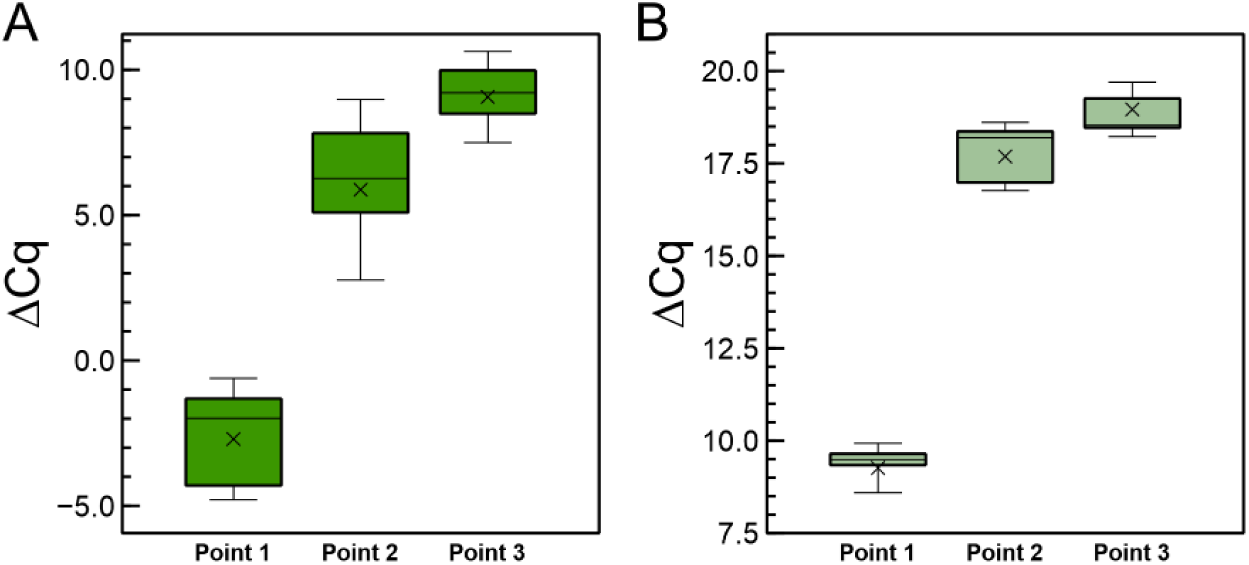
Quantitative differences of the CP-(ds)RNA (A) and the 6125-vsiRNA (B) detected by RT-qPCR at 3 dpt after spraying the CP-dsRNA in local (point 1), close distal (point 2) and far distal (point 3) sampling as described in Fig. 8.

When RNAi was investigated, we found that the 6125-vsiRNAs could be detected in non-treated leaves at 3 dpt, indicating systemic long-distance movement of the silencing signal and confirming the earlier experiments (Fig. 10B). Comparison of ΔCq_6125-vsiRNAs-5.8S siRNA_ between the point of spraying (point 1) and points 2 and 3 showed that in the former, 9.1 × 10^3^ -fold and 3.3 × 10^4^ -times, respectively, more vsiRNA was detected. In point 3 (far distal), 7-fold less 6125-vsiRNA was detected in comparison with point 2 (close distal). Thus, the dsRNA/vsiRNAs reaching distal sites were very diluted in comparison to amounts quantified in the point of application. When we compared the ratios between long (ds)RNAs and vsiRNAs in the different sampling sites, it resulted that they were similar in points 1 and 2, but when compared between point 1 and point 3, the relative ratio was 7.9 times higher for the vsiRNAs in the latter. Therefore, although some correlation was found in the ratios, the comparatively higher ratio of vsiRNAs in point 3 (opposite leaf), might suggest a local multiplication of vsiRNAs or different phloem systemic movement of long (ds)RNAs depending on the location.

Finally, the stability of the (ds)RNA in the leaves was also investigated. After the application of the CP-dsRNAs on the leaves, we washed them and took samples from the leaf sprayed at 3, 6 and 10 dpt, for RNA extractions and RT-qPCR. The result showed that the dsRNA could be detected at least up to 10 days following the treatments (Fig. 11). Interestingly, the amount of (ds)RNA was almost stable between 6 and 10 dpt at the point of application. Another sampling was done at the apex and the (ds)RNA could be detected as well. It was diluted 7.9 × 10^5^-fold with respect to amount quantified in the sprayed leaf, similarly to the ratios found at 3 dpt between the leaf sprayed and the opposite leaf.

**Fig. 11.**
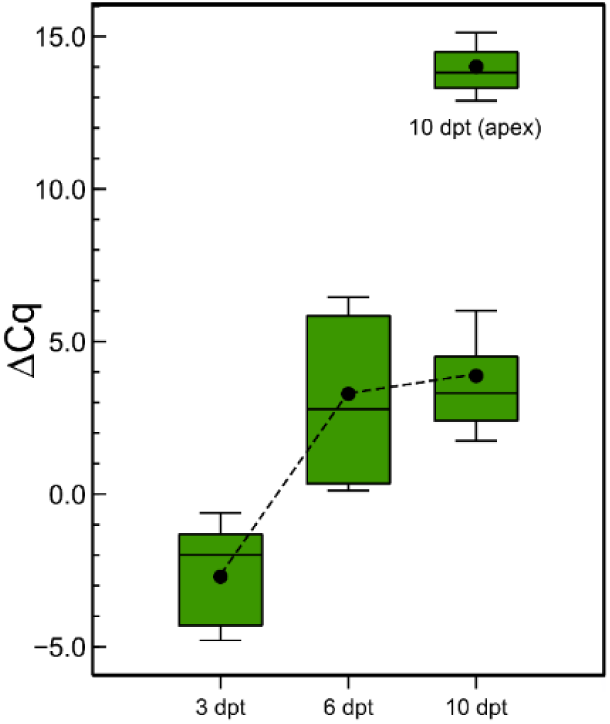
Stability of the dsRNA in the plants after topical application. Detection by RT-qPCR of cp- (ds)RNA at 3, 6 and 10 days post CP-dsRNA application using the spraying. Another sampling was done at the apex at 10 dpt.

### 3.7. Failure in eliciting protection by exogenous dsRNAs in ToLCNDV infections

To investigate if a similar effect of exogenously applied dsRNAs protects zucchini against ToLCNDV, we carried out parallel experiments to those performed for CGMMV. Groups of eight plants were either sprayed with AV1/BC1-dsRNAs or agroinfiltrated with the pGHE-AV1 and pGHE-BC1 constructions. They were infected at 3 dpt with viruliferous insects hosting ToLCNDV-ES and kept in insect proof cages in the growth chamber. Another group of plants was mock-treated for comparisons. At 15 dpi we observed no differences in the number of symptomatic plants nor in the severity of symptoms between the treated and the untreated plants, either sprayed with dsRNA or agroinoculated.

## 4 Discussion

Control of CGMMV by RNAi has been achieved in *Nicotiana benthamiana* using a transgenic approach with a transgene designed to produce an inverted repeat RNA containing the *cp* gene separated by an intron (Kamachi et al., 2007). In watermelon, a single chimeric transgene comprising a silencer DNA from the partial *N* gene of watermelon silver mottle virus (WSMoV) fused to the partial *cp* gene sequences of cucumber mosaic virus (CMV), watermelon mosaic virus (WMV) and CGMMV conferred multiple virus resistance (Lin et al., 2012). More recently, Miao et al. (2021) have designed polycistronic artificial microRNAs that, when externally applied by a transient expression system, limit CGMMV disease in cucumber and *N. benthamiana*. Transgenic cucumber lines expressing the polycistronic artificial microRNAs also developed resistance to the virus.

In our approach, direct application of dsRNAs was aimed to control CGMMV disease in cucumber under greenhouse conditions. Transient expression of dsRNA triggered by a plasmid construction with direct and inverted repeats of the *cp* and *mp* genes of CGMMV were also capable of limiting disease development in cucumber during agroinoculation experiments performed in a phytotron. The effect of dsRNA application in limiting CGMMV disease progress and virus accumulation resulted more consistent in trials performed during spring than in summer. Higher temperatures limit plants tolerance to this virus (Dombrovsky et al., 2017) and increase its mobility in the plant (Moreno et al., 2004). Nevertheless, a reduction in virus accumulation in the plants will help limiting the damages and will probably reduce the transmission between plants during in routine crop management practices in the greenhouse such as planting, pruning, harvesting, and others. The reduction in virus accumulation in greenhouse conditions of 48- and 5-fold depending on the season was comparable to the results obtained in groundnut bud necrosis virus (GBNV), where it has been reported 20- and 12.5-fold reduction after rubbing dsRNA on plant leaves in cowpea and *N. benthamiana*, respectively (Gupta et al., 2021). The correlation between symptom severity and virus accumulation has been previously reported for many plant pathosystems, specifically for CGMMV in *N. benthamiana* by ELISA (Ali et al., 2016), in cucumber by RT-qPCR (Crespo et al., 2018), and for CVYV in cucumber, also by RT-qPCR (Galipienso et al., 2013). The introduction of resistance in transgenic plants expressing virus genes leads to reduced symptom expression that is correlated with a decrease in virus accumulation (Bagewadi et al., 2014). Similarly, induced resistance with exogenous delivery of dsRNA or siRNAs leads to reduction in virus accumulation in plants as shown by serology or qPCR (Tenllado et al., 2003; Mitter et al., 2017a; Rego-Machado et al., 2020; Holeva et al., 2021; Gupta et al., 2021; Necira et al., 2021).

Plant cuticles are barriers to dsRNA entry that has also to reach the cell wall and cross the plasma membrane. High pressure spraying allows the entry of the dsRNA, as well as do lesions made by abrasives such as carborundum, like in our experiments when applied the dsRNA by rubbing or celite, and as reported for groundnut bud necrosis virus (GBNV) control using dsRNA (Gupta et al., 2021). Other authors have proposed surfactants to facilitate the entry of the dsRNA and the movement of the silencing signals (Schwartz et al., 2020; Bennet et al., 2021). However, in the case of CGMMV control in cucumber using RNAi, we believe that surfactants should be avoided, because in preliminary assays we have observed that disease severity increased and the time to symptom onset shortened when the surfactant BREAK-THRU S279 (BASF) was included in dsRNA formulations for CGMMV control (results not shown). Formulations that include nanoparticles to protect the si/dsRNAs from degradation and facilitate the entry to the plant seem promising (Uslu and Wassenegger, 2020). Therefore, there is room for improvement in dsRNA applications, given the variability in plant responsiveness to dsRNAs for CGMMV protection that we have observed. This can be due to differences in dosage, pressure applied, phenological state of the plant, etc. A high pressure of 7-8 bars seems to be necessary for triggering RNAi responses in GFP transgenic *N. benthamiana* 16c (Dalakouras et al., 2016); in our case 2.5 bars resulted to be sufficient to induce CGMMV resistance in cucumber.

By densitometry analysis, we previously estimated the amount of CP-dsRNA present in the RNA extractions from the *E. coli* cells to be 62 ng dsRNA/μg total RNA (Delgado and Velasco, 2021). Thus, when spraying 120 μg of the bacterial RNA extraction, that include ssRNA and DNA from 5 mL of bacterial culture, the amount of dsRNA applied on the plants, that was effective for limiting CGMMV disease, was 3.7 μg in total, or 35 ng·cm^-2^ in each of the two leaves/plant on average. In tomato, the control of the tomato mosaic virus (ToMV) has been achieved using 200-400 μg of bacterial dsRNA extract, obtained using a similar method (Rego-Machado et al., 2020). Gupta et al. (2020) used 5 μg of dsRNA/plant for limiting GBNV disease in *N. benthamiana*, but in this case, the dsRNA extract was treated with DNase I and RNAse A to remove the DNA and ssRNA, so this quantity cannot be directly comparable to ours. In another example, 1.25 μg of *in vitro* transcribed dsRNA was effective for the control of pepper mild mottle virus (PMMoV) in *N. tabacum* (Mitter et al. 2017a). For inducing resistance to tobacco mosaic virus (TMV) in tobacco, 300 μg of crude bacterially expressed dsRNA per plant were used (Yin et al., 2009). The control of papaya ringspot virus (PRSV) in papaya was achieved using 100 μg/plant of bacterially expressed dsRNA (Shen et al., 2014). In general, lower amounts have been used in the case of *in vitro* synthesized dsRNA (reviewed in: Dubrovina and Kiselev, 2019). For example, for the protection of zucchini against Zucchini yellow mosaic virus (ZYMV), 60 μg/plant of *in vitro* synthesized dsRNA has been applied (Kaldis et al., 2018). Thus, there is a lack of homogeneity in the description of quantity and conditions of exogenous dsRNA applications, making it difficult to compare the amount of effective dsRNA applied as in the literature each author reports different synthesis approaches (*in vitro, in vivo*), extractions methods, preliminary treatments with DNase and/or RNAse A, etc. (Dubrovina and Kiselev, 2019; Das et al., 2020). We consider that the net volume of bacterial culture per plant, which in our case for CGMMV treatments was 5 mL/plant that can be used to compare the different treatments so far reported that include *in vitro* or *in vivo* dsRNA synthesis.

Analysis of vsiRNAs in CGMMV infected plants have been reported previously. In *Lagenaria siceraria*, the sRNAs sizing 24-nt length were predominant over the 21, 22 or 23-nt in leaves or fruits (Li et al., 2016a). Similarly, in cucumber, other authors reported the 24-nt sRNAs as predominant in CGMMV infected or healthy plants (Li et al., 2016b). Other authors found that the 23-nt class of sRNAs was predominant in CGMMV infected cucumber (Liu et al., 2015), in contrast to what was reported in healthy cucumber plants, where the 24-nt class of sRNAs was predominant (Martinez et al., 2015). On the contrary, in cucumber we observed the 21-nt followed by the 22-nt as the predominant sRNAs in leaves. Regarding the specific CGMMV vsiRNAs, we as well as other authors observed the 21-nt class as predominant, followed by the 22-nt class. In consequence, the involvement of the DICER-LIKE endonucleases DCL4 and DCL2 seem to be predominant in the processing of the exogeneous CP-dsRNA (Liu et al., 2009). In contrast to the previous report in cucumber and *L. siceraria*, we observed that the negative CGMMV vsiRNAs sequences were predominant in the leaves. With respect to the preferred 5’ and 3’ terminal nucleotides of the vsiRNAs, we observed that both in dsRNA treated or untreated plants, the 5’-A and the 3’-U were predominant, confirming what has been reported for vsiRNAs of plants viruses, that usually start with A or U (Donaire et al., 2009). However, Li et al. (2016b) reported a predominance of a 5’-C, followed by 5’-A in CGMMV vsiRNAs in cucumber. In *L. siceraria*, the 5’-A was reported as the CGMMV predominant nucleotide end of the vsiRNAs in fruits and 5’-U in leaves (Li et al, 2016a). In ToMV in tomato, the 5’-U terminal nucleotide in the 21-nt class was predominant (Riego-Machado et al., 2020) in dsRNA-treated and non-treated plants. Given that AGO1 and AGO2 preferentially recruit small RNAs with a 5′ terminal of U and A, respectively, it seems that several ARGONAUTE proteins are involved in the processing of the vsiRNAs within the RISC complex in cucumber (Mi et al., 2008). Hotspots of CGMMV-derived vsiRNAs could be identified in dsRNA-treated and not treated plants, that were predominant in the 5’ and 3’ end of the viral genome. The distribution of the vsiRNAs was similar in both cases, although there was a bias for 5’ hotspots in the non-dsRNA treated plants. In CGMMV infected cucumber, hotspots were already reported in the terminal ends of the genome (Li et al, 2016b). This is similar to the findings in *L. siceraria* fruits, but not in leaves, where the distribution of vsiRNA hotspots showed no predominance along the genome (Li et al, 2016a). The proportion vsiRNA classes was similar in dsRNA treated or untreated plants as reported before (Riego-Machado et al., 2020); however, we observed a 40% reduction in the number of vsiRNAs of all the classes in the dsRNA-treated plants. The *cp* gene of CGMMV is expressed from subgenomic RNAs which might explain the presence of abundant vsiRNA hotspots in this region (Adams et al., 2017). Besides, we found that many vsiRNAs were produced in the 3′ untranslated region, that conforms a tRNA-like structure.

In this work, we report that long specific RNAs can be detected systemically in distal part of the plant after the application of the dsRNAs, either by spraying or elicited by agroinfiltration, and are effective in limiting CGMMV accumulation. This could also be obtained by using LDH-nanoparticles that released gradually the dsRNA on the leaves of plants (Mitter et al., 2017a). However, in our case, the release of dsRNA encapsulated in layered double hydroxides (LDH) nanoparticles does not improve CGMMV resistance with respect to naked dsRNA (manuscript in preparation). Detection of dsRNAs was possible by RT-qPCR at the site of spraying and in proximal half of the leaf at 3 dpt, showing long-distance transport of the dsRNA. There is strong evidence for systemic movement of siRNAs to short distances (through a few cells) throughout the plasmodesmata without producing secondary siRNAs (Kim, 2005). However, long-distance movement of sRNAs and systemic silencing seems to require the amplification of the silencing signals by RdRps in the phloem (Dalakouras et al., 2018). Thus, the mechanism seems to involve long distance movements of the dsRNA in the apoplast and the phloem and translocation from there to the symplast for eliciting RNAi and subsequent siRNA production (Das et al., 2020). Recently, when applying exogenous ZYMV-derived dsRNAs, long-distance transport of long dsRNAs has been shown by semiquantitive RT-PCR even at 21 dpt (Kaldis et al., 2018). We detected vsiRNAs following dsRNA application, at the site of application, as well as in the distal part of the leaf and in another opposite leaf, suggesting that the vsiRNAs either make long-distance movements or are the result of local RNAi processing of the moving (ds)RNAs. Local and systemic vsiRNAs have been identified from ZYMV-derived vsiRNAs after dsRNA rubbing (Kaldis et al., 2018). In another report, ToMV-derived dsRNA was detected by RT-PCR at 10 dpt at the site of application but could not be detected in distal leaves (Rego-Machado et al., 2020); the dsRNA-derived vsiRNAs were identified by HTS in local and distal leaves of the application site. A PMMoV-derived dsRNA was detected by Northern-blot in pepper at 7 dpt (Tenllado and Díaz-Ruiz, 2001). More recently, a GBNV-derived dsRNA was locally and systemically detected by semiquantitative RT-qPCR at 7 dpt in *N. benthamiana* and at 5 dpt in cowpea (Gupta et al, 2021). Delivery of insect-derived dsRNA into barley plants showed systemic dsRNA movement and distal siRNAs were present as shown by HTS (Biedenkopf et al., 2020). Conceivably, using the RT-qPCR, more sensitive, than the conventional RT-PCR allowed us to detect systemic movements of the (ds)RNAs, that were several magnitude orders more diluted with respect to the site of application. Therefore, our work supports the evidence for systemic movement of long RNA molecules after exogenous application. Moreover, we have shown a correlation between the amounts of dsRNA, either applied or systemic, and derived siRNAs. On the other hand, we propose a method for the direct quantitation of (ds)RNA in plants after the application and washing up the surface, that yields a comparison test for procedures and formulations for dsRNA delivery. The protection effect of naked dsRNA is probably limited to shorts periods after its application, although the dsRNA could be detected in the site of application at least up to 9 dpt (Kaldis et al., 2018) or even at 10 dpt, as we have observed. This is a serious limitation in the management of the diseases, so that either repeated dsRNA applications are performed, or the dsRNA applied in the first instance must endure for a longer period. Mitter et al. (2017a) elongated that availability of dsRNA in the plants up to 20 dpt by the controlled release on the leaf surface mediated by nanoparticles. Thus, it is plausible that improving the administration of dsRNAs will enhance their efficacy in increasing resistance to viruses.

DsRNA exogenously applied offers a promising tool for virus control, although it might not be effective in all cases, as is described for DNA viruses, such as the begomoviruses tomato severe rugose virus (ToSRV) in tomato (Rego-Machado et al., 2020), or ToLCNDV in zucchini squash, in the present work. Nevertheless, there is potential for improvement in the case of DNA viruses, as transgenic plants expressing microRNAs specific for ToLCNDV increased virus tolerance in tomato (Van Vu et al., 2013). Conceivably, increasing the amount of the dsRNA applied or improving the formulations may contribute to find effective exogenous dsRNA application in those viruses. Finally, we have shown that cucumber plants topically treated with dsRNA limit the multiplication of CGMMV and, consequently, the expression of symptoms and other deleterious effects of the disease. We postulate that it may be necessary to maintain the uptake or continuous application of dsRNA to the plant during the vegetative period, particularly in summer, either by increasing the number of doses or by generating a controlled release by encapsulation of dsRNA in nanocomposites, in order to make the treatments more reliable in field applications (Rank and Koch., 2021).

## 6 Author Contributions

L.V. and J.D. designed the research; J.D., L.R., D.J. and L.V. performed the experiments; J.D., L.R., D.J. and L. V. analysed the data and results; L.V. wrote the original draft; D.J., J.D., L.R. and L.V. wrote and edited the final manuscript; L.V. and D.J. provided funds for the project.

## 8 Acknowledgments

The authors acknowledge E. Martínez-Campos, A. Delgado-Olidén and A. Belmonte for the technical assistance.

## 10 Supplementary Material

Supplementary Fig. 1. Rating scale used to score symptom severity in CGMMV cucumber plants.

Supplementary Fig. 2. Agarose gel (2%) of total RNA extracts from IPTG-induced HT115(DE3) cells.

Supplementary Fig. 3. Unique (upper) and total (lower) read length distribution of the small RNAs for the MO (left) and DS (right) samples.

Supplementary Fig. 4. Small RNA distribution as function of read length and type for A) DS sample (pool of CP-dsRNA treated plants) and B) MO sample (pool of untreated plants).

Supplementary Fig. 5. Proportion of vsiRNAs according to their length aligning to the CGMMV genome in samples MO and DS.

Supplementary Fig. 6. Nucleotide preferences at 5’ and 3’ terminal ends of the CGMMV vsiRNAs from A) sample MO and B) sample DS.

Supplementary Fig. 7. Melting curves for the amplification of the vsiRNAs.

Supplementary Fig. 8. Systemic movement of (ds)RNAs and vsiRNAs was evaluated in agroinoculated plants.

Supplementary Fig. 9. Systemic movement of sprayed dsRNA.

Supplementary Table 1. Primers used in this work.

Supplementary. Table 2. Primers used in this work for obtaining the plasmids pGHE-AV1, pGHE-BC1, pGHE-CP and pGHE-MP used in the agroinoculations.

## 11 Data Availability Statement

Data available on request from the authors.

