## Supplementary material for "Exogenous application of dsRNA for the control of viruses in cucurbits"

**SUPPLEMENTARY MATERIALS**

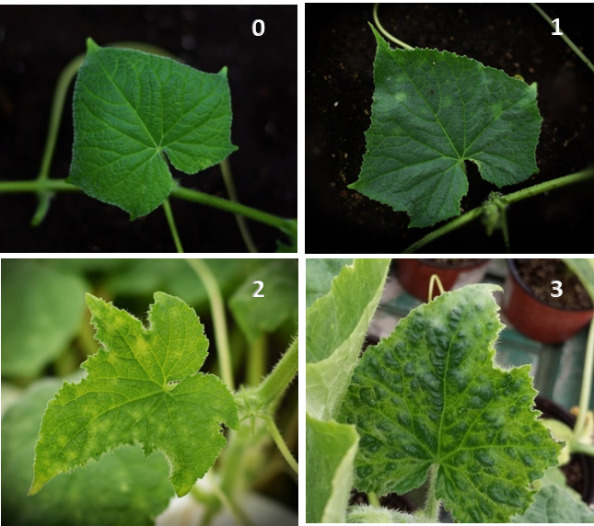

**Supp. Fig. 1.** Rating scale used to score symptom severity in CGMMV cucumber plants: (0) asymptomatic; (1) mild symptoms, (2) severe symptoms and (3) very severe symptoms.

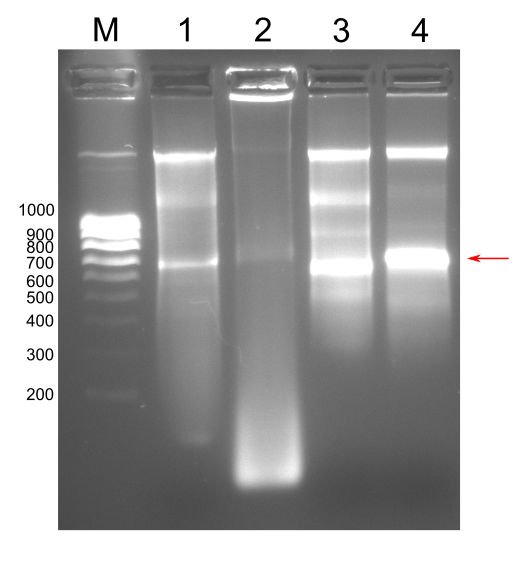

**Supp. Fig. 2.** Agarose gel (2%) of total RNA extracts from IPTG-induced HT115(DE3) cells harboring (1) L4440-AV1, (2) L4440-BC1, (3) L4440-CP and (4) L4440-MP. DsRNA bands are indicated with an arrow. M: molecular weight marker NZYDNA Ladder V.

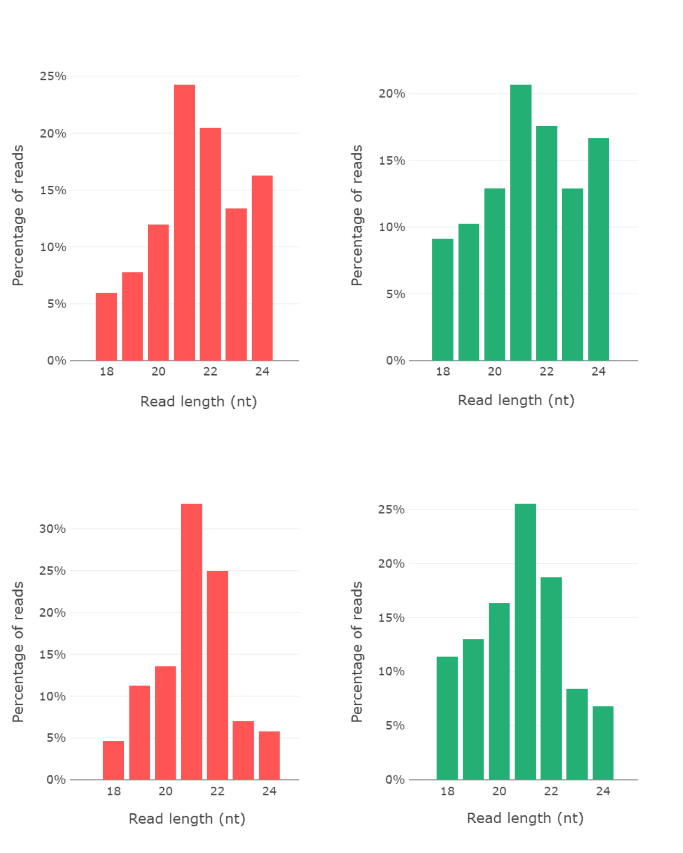

**Supp. Fig. 3**. Unique (upper) and total (lower) read length distribution of the small RNAs for the MO (left) and DS (right) samples.

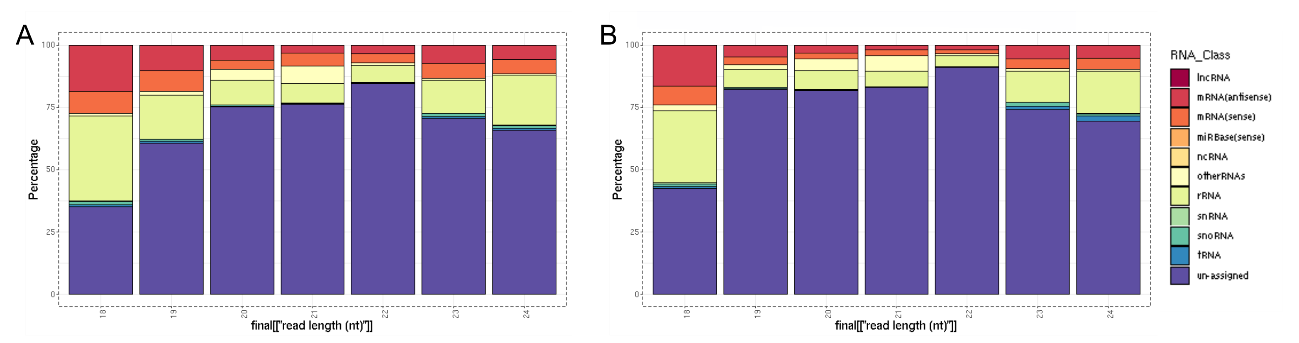

**Supp. Fig. 4**. Small RNA distribution as function of read length and type for A) DS sample (pool of CP-dsRNA treated plants) and B) MO sample (pool of untreated plants).

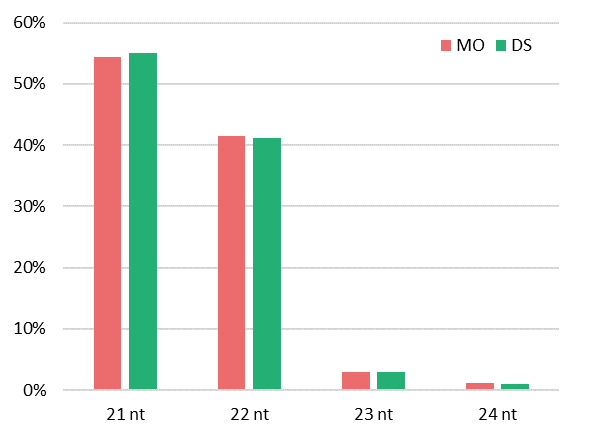

**Supp. Fig. 5**. Proportion of vsiRNAs according to their length aligning to the CGMMV genome in samples MO and DS.

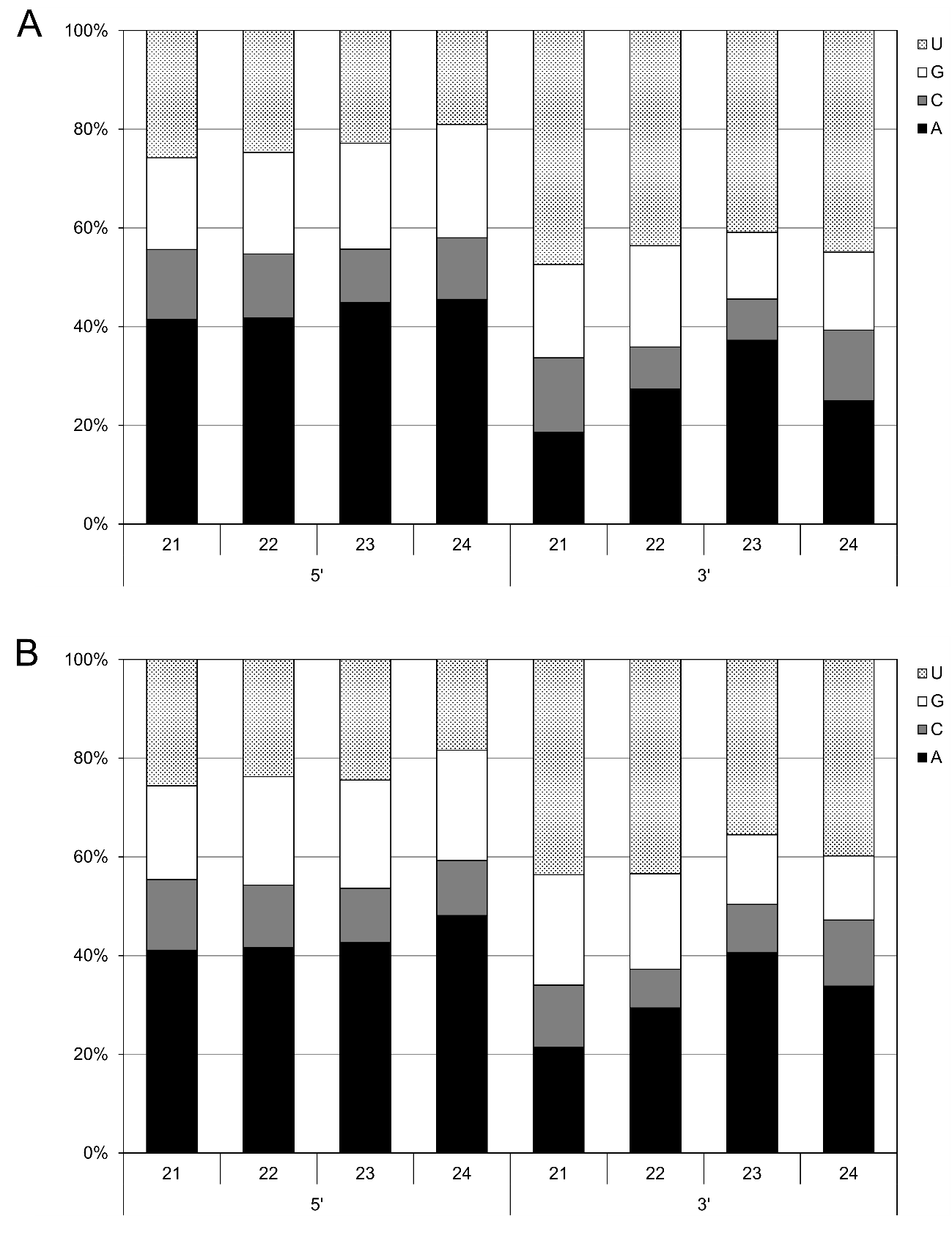

**Supp. Fig. 6**. Nucleotide preferences at 5’ and 3’ terminal ends of the CGMMV vsiRNAs from A) sample MO and B) sample DS.

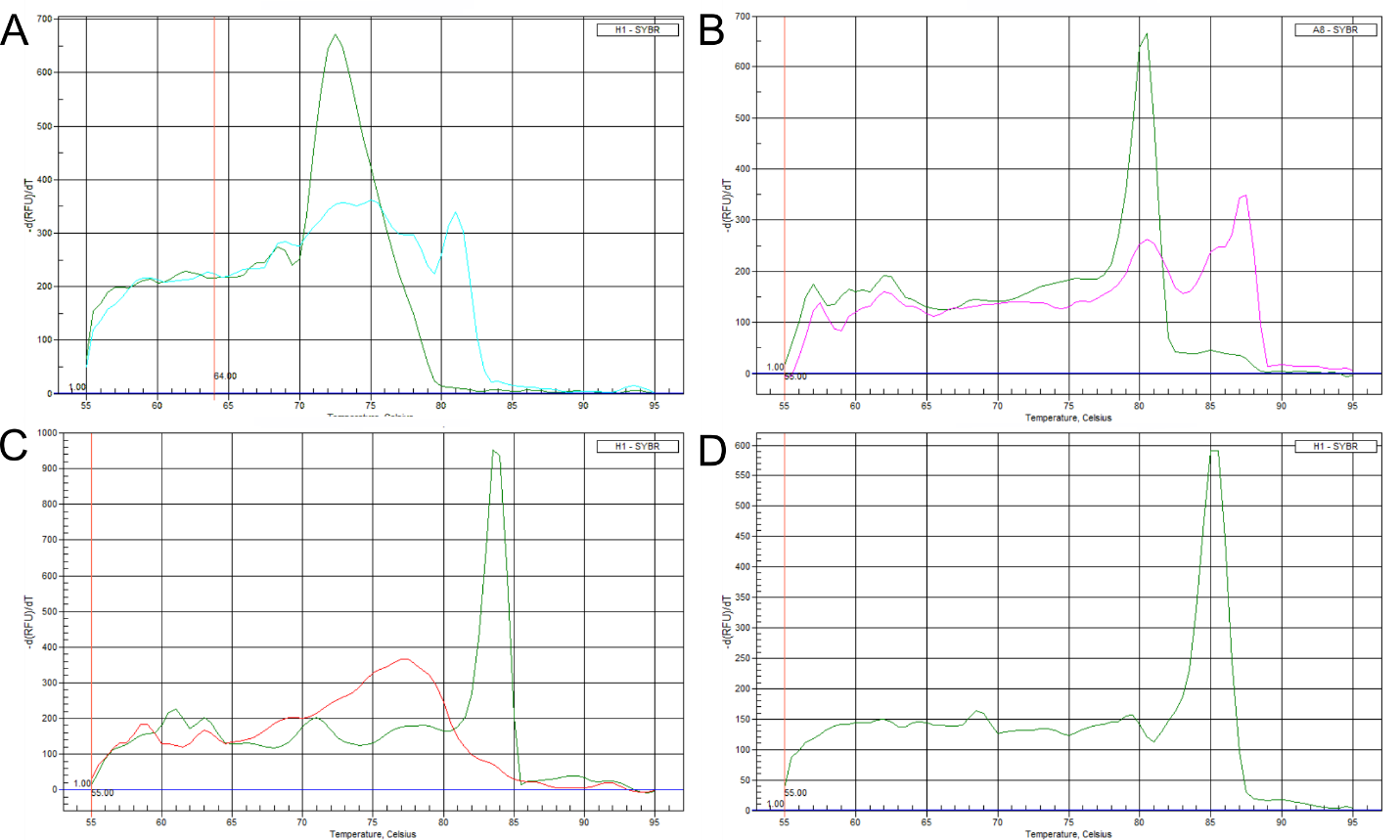

**Supp. Fig. 7**. Melting curves for the amplification of the vsiRNAs: A) 1570-vsiRNA, B) 5234-vsiRNA, C) 6125-vsiRNA and D) the small rRNA-5.8S used as a reference. Green lanes correspond to the positive samples and blue, red or pink lanes correspond to the negative ones.

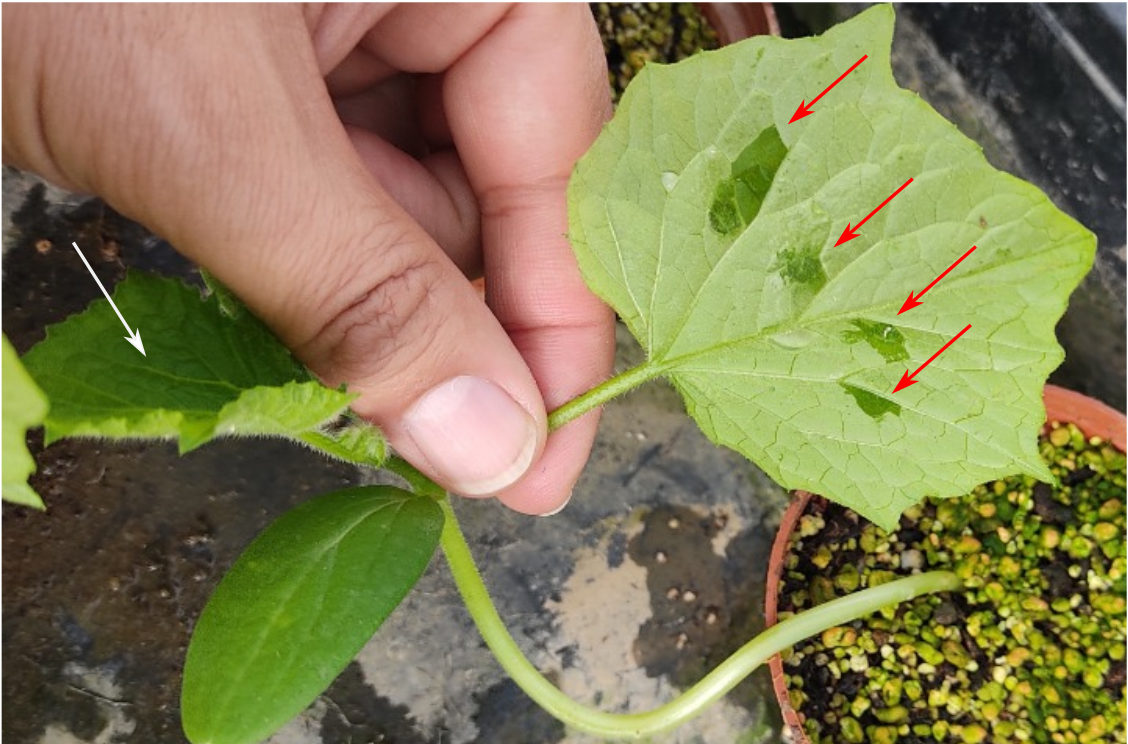

**Supp. Fig. 8**. Systemic movement of (ds)RNAs and vsiRNAs was evaluated in agroinoculated plants. Leaves of cucumber plants were agroinoculated for dsRNA expression (red arrows) in four points (50 μL of bacterial inoculant each). Three days after the agroinoculation, samples were taken from the distal leaves indicated with the white arrow (left). These samples were used for long RNA and vsiRNA detection.

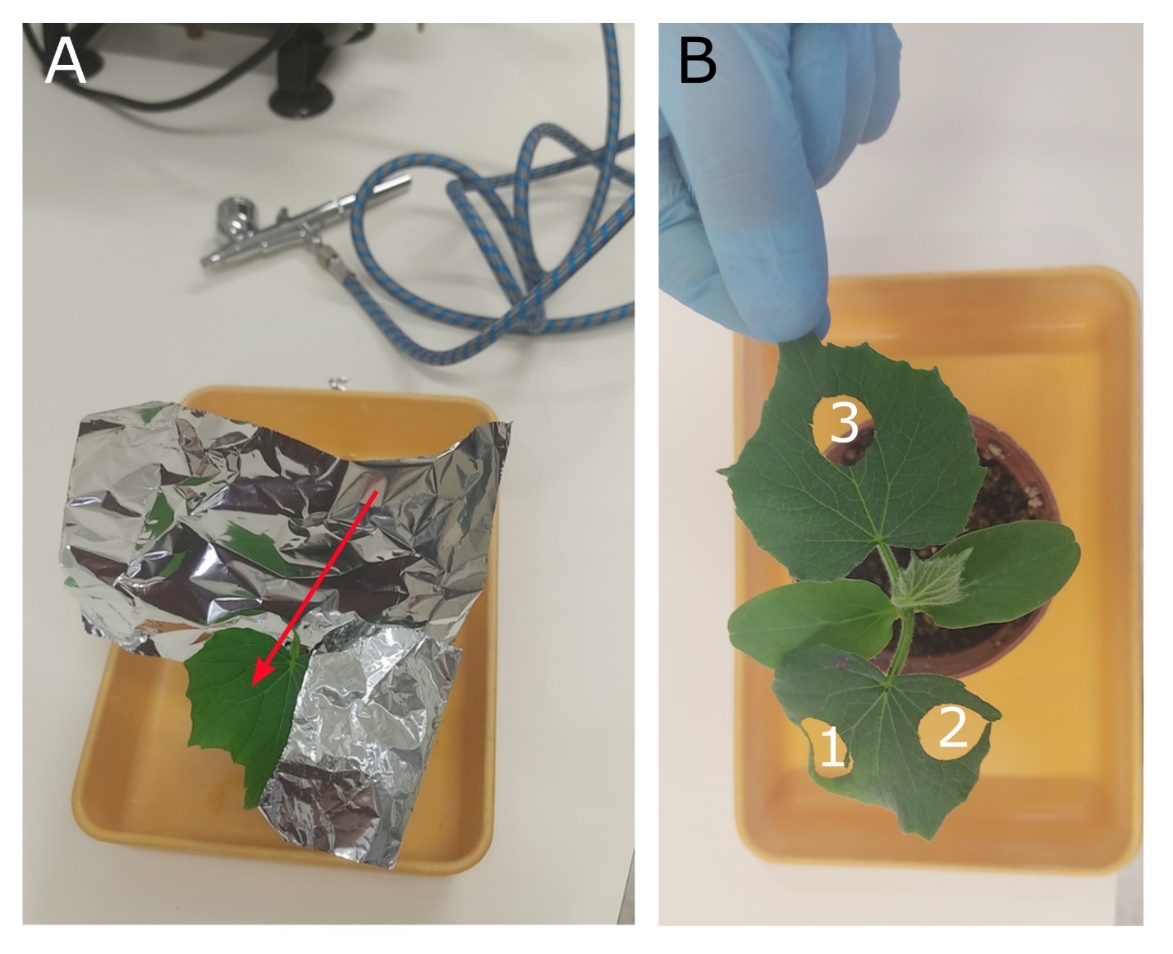

**Supp. Fig. 9.** Systemic movement of sprayed dsRNA. The point of application of the dsRNA is indicated with an arrow. Half of the leaf and the rest of the plant is foil-covered prior application (A) followed by washing with distilled water. The points of sampling for the analyses are shown and numbered in (B).

**Supp. Table 1.** Primers used in this work.

| **Name** | **Target** | **Sequence 5’-3’** | **Source** |
| --- | --- | --- | --- |
| attB1n-AV1F | AV1 ToLCNDV | GGGGACAAGTTTGTACAAAAAAGCAGGCTTATCTCCAGACACAGTCGGCTA | This work |
| attB2n-AV1R | “ | GGGGACCACTTTGTACAAGAAAGCTGGGTAATGTTCTTCACCGTGGCTGT | “ |
| attB1n-BC1F | BC1 ToLCNDV | GGGGACAAGTTTGTACAAAAAAGCAGGCTTACTCCGATCTGCTAGCCCATG | “ |
| attB2n-BC1R | “ | GGGGACCACTTTGTACAAGAAAGCTGGGTAACGATGCTGCAGAGGTAACC | “ |
| attB1n-CPF | CP CGMMV | GGGGACAAGTTTGTACAAAAAAGCAGGCTTACAATCCGATCACACCTAGCA | Delgado and Velasco, 2021 |
| attB2n-CPR | “ | GGGGACCACTTTGTACAAGAAAGCTGGGTATGGTAGCCTCTGACCAGACT | “ |
| attB1n-MPF | MP CGMMV | GGGGACAAGTTTGTACAAAAAAGCAGGCTTACCTGTCAAGTTGTTGCGTGG | This work |
| attB2n-MPR | “ | GGGGACCACTTTGTACAAGAAAGCTGGGTAACAGACTGCGACCTAGACCT | “ |
| CP197F | Cp CGMMV | TACGCTTTCCTCAACGGTCC | Delgado and Velasco, 2021 |
| CP305R | “ | GCGTCGGATTGCTAGGATCT | “ |
| QMP1F | Mp CGMMV | CCTGTCAAGTTGTTGCGTGG | This work |
| QMP110R | “ | GCACCACCCCTACAGGATTC | “ |
| CUC18S-For | 18S rRNA *C. sativus* | GGCGGATGTTGCTTTAAGGA | Gil-Salas et al., 2009 |
| CUC18S-Rev | “ | GTGGTGCCCTTCCGTCAAT | “ |
| CG-1570 | vsiRNA-1570 (RdRp) | AAGTATTGCTACCTGAACCGTT | This work |
| CG-6125 | vsiRNA-6125 (Cp) | GCTAGGGCTGAGATAGATAATT | “ |
| CG-5234 | vsiRNA-5234 (Mp) | GGAACGTACCGGAATCCTGTA | “ |
| PolyT | Universal polyT primer | GCGAGCACAGAATTAATACGACTCACTATAGGTTTTTTTTTTTTVN | Shi and Chiang, 2005 |
| URP | Universal reverse primer | GCGAGCACAGAATTAATACGAC | “ |
| CUC5.8S | 5.8S sRNA *C. sativus* | CTTGGTGTGAATTGCAGGATC | This work |

**Supp. Table 2.** Primers used in this work for obtaining the plasmids pGHE-AV1, pGHE-BC1, pGHE-CP and pGHE-MP used in the agroinoculations.

| **Name** | **Sequence (5’-3’)** | **Function** |
| --- | --- | --- |
| CP-GIB1 | CATTTGGAGAGGACACGCGTACAAAAAAGCAGGCTCAATCC | Gibson assembly of the CP gene from CGMMV to pHellsgate8 |
| CP-GIB2 | TTGGATCCTAAAGCTGGGTTGGTAGCCTC |  |
| CP-GIB3 | CCGAATTCCAAAGCTGGGTTGGTAGCCTC |  |
| CP-GIB4 | TCTCATTAAAGCAGGACTGTACAAAAAAGCAGGCTCAATCC |  |
| CP-GIB5 | CCCAGCTTTGGAATTCGGTACCCCAGCTT |  |
| CP-GIB6 | CCCAGCTTTAGGATCCAAGCTTATCGATTTCGA |  |
| MP1-GIB | CATTTGGAGAGGACACGCGCTCCTGTCAAGTTGTTGCG | Gibson assembly of the MP gene from CGMMV to pHellsgate8 |
| MP2-GIB | CCGAATTCCTACAGACTGCGACCTAGACCT |  |
| MP3-GIB | CAGTCTGTAGGAATTCGGTACCCCAGCTT |  |
| MP4-GIB | CAGTCTGTAAGGATCCAAGCTTATCGATTTCGA |  |
| MP5-GIB | TTGGATCCTTACAGACTGCGACCTAGACCT |  |
| MP6-GIB | TCTCATTAAAGCAGGACTGCTCCTGTCAAGTTGTTGCG |  |
| AV-GIB1 | CATTTGGAGAGGACACGCGCTTCTCCAGACACAGTCGG | Gibson assembly of the AV1 gene from ToLCNDV to pHellsgate8 |
| AV-GIB2 | CCGAATTCCGGTATGTTCTTCACCGTGGC |  |
| AV-GIB3 | GAACATACCGGAATTCGGTACCCCAGCTT |  |
| AV-GIB4 | GAACATACCAGGATCCAAGCTTATCGATTTCGA |  |
| AV-GIB5 | TTGGATCCTGGTATGTTCTTCACCGTGGC |  |
| AV-GIB6 | TCTCATTAAAGCAGGACTGCTTCTCCAGACACAGTCGG |  |
| BC1-GIB | CATTTGGAGAGGACACGCCTCCGATCTGCTAGCCCATG | Gibson assembly of the BC1 gene from ToLCNDV to pHellsgate8 |
| BC2-GIB | CCGAATTCCATGTCATTTCCTTCCATGTTCGA |  |
| BC3-GIB | AAATGACATGGAATTCGGTACCCCAGCTT |  |
| BC4-GIB | GAAATGACATAGGATCCAAGCTTATCGATTTCGA |  |
| BC5-GIB | CTTGGATCCTATGTCATTTCCTTCCATGTTCGA |  |
| BC6-GIB | TCTCATTAAAGCAGGACTCTCCGATCTGCTAGCCCATG |  |

**Supp. Table 3.** Linear regression analysis of length vs. dry weight of the plants in the experiments carried out in spring and summer among the different conditions.

| **Predictor ^a^** | **Estimate** | **SE** | **t** | ***P*** |
| --- | --- | --- | --- | --- |
| **Spring assay** |  |  |  |  |
| Intercept ^b^ | 48.367 | 3.469 | 13.944 | < .001 |
| Dry weight (g) | 6.084 | 0.603 | 10.097 | < .001 |
| Condition: |  |  |  |  |
| dRNA-treated – Non-infected | 7.282 | 2.152 | 3.384 | 0.002 |
| untreated – Non-infected | -0.353 | 2.335 | -0.151 | 0.881 |
| **Summer assay** |  |  |  |  |
| Intercept ^b^ | 147.25 | 14.29 | 10.31 | < .001 |
| Dry weight (g) | 8.26 | 1.06 | 7.78 | < .001 |
| Condition: |  |  |  |  |
| dRNA-treated – Non-infected | -18.37 | 11.13 | -1.65 | 0.107 |
| untreated – Non-infected | -51.89 | 11.12 | -4.67 | < .001 |
| ᵃ Model coefficients: Length; ^b^ Represents reference level |  |  |  |  |
